# A Drug Repurposing Approach Reveals Targetable Epigenetic Pathways in *Plasmodium vivax* Hypnozoites

**DOI:** 10.1101/2023.01.31.526483

**Authors:** S. P. Maher, M. A. Bakowski, A. Vantaux, E. L. Flannery, C. Andolina, M. Gupta, Y. Antonova-Koch, M. Argomaniz, M. Cabrera-Mora, B. Campo, A. T. Chao, A. K. Chatterjee, W. T. Cheng, E. Chuenchob, C. A. Cooper, K. Cottier, M. R. Galinski, A. Harupa-Chung, H. Ji, S. B. Joseph, T. Lenz, S. Lonardi, J. Matheson, S. A. Mikolajczak, T. Moeller, A. Orban, V. Padín-Irizarry, K. Pan, J. Péneau, J. Prudhomme, C. Roesch, A. A. Ruberto, S. S. Sabnis, C. L. Saney, J. Sattabongkot, S. Sereshki, S. Suriyakan, R. Ubalee, Y. Wang, P. Wasisakun, J. Yin, J. Popovici, C. W. McNamara, C. J. Joyner, F. Nosten, B. Witkowski, K. G. Le Roch, D. E. Kyle

## Abstract

Radical cure of *Plasmodium vivax* malaria must include elimination of quiescent ‘hypnozoite’ forms in the liver; however, the only FDA-approved treatments are contraindicated in many vulnerable populations. To identify new drugs and drug targets for hypnozoites, we screened the Repurposing, Focused Rescue, and Accelerated Medchem (ReFRAME) library and a collection of epigenetic inhibitors against *P. vivax* liver stages. From both libraries, we identified inhibitors targeting epigenetics pathways as selectively active against *P. vivax* and *P. cynomolgi* hypnozoites. These include DNA methyltransferase (DNMT) inhibitors as well as several inhibitors targeting histone post-translational modifications. Immunofluorescence staining of *Plasmodium* liver forms showed strong nuclear 5-methylcystosine signal, indicating liver stage parasite DNA is methylated. Using bisulfite sequencing, we mapped genomic DNA methylation in sporozoites, revealing DNA methylation signals in most coding genes. We also demonstrated that methylation level in proximal promoter regions as well as in the first exon of the genes may affect, at least partially, gene expression in *P. vivax*. The importance of selective inhibitors targeting epigenetic features on hypnozoites was validated using MMV019721, an acetyl-CoA synthetase inhibitor that affects histone acetylation and was previously reported as active against *P. falciparum* blood stages. In summary, our data indicate that several epigenetic mechanisms are likely modulating hypnozoite formation or persistence and provide an avenue for the discovery and development of improved radical cure antimalarials.

**One-Sentence Summary:** Drug repurposing screens reveal several epigenetic inhibitors as active against *P. vivax* hypnozoites demonstrating that epigenetic pathways play a central role in hypnozoite quiescence.

## Introduction

Of the six species of *Plasmodium* that cause malaria in humans^1^, *Plasmodium vivax* is the most globally widespread^2^. Vivax malaria now accounts for the most malaria episodes in countries with successful falciparum malaria control programs^3^. Controlling vivax malaria is complicated by the ability of *P. vivax* sporozoites, the infectious stage inoculated by mosquitoes, to invade hepatocytes and become quiescent^4,5^. These quiescent ‘hypnozoites’ persist, undetectable, for months or even years before resuming growth and initiating a ‘relapse’ blood stage infection, leading to subsequent transmission back to mosquitoes^6^. New evidence suggests this transmission is expedited and silent as *P. vivax* liver merozoites can immediately form gametocytes instead of first having to establish an asexual stage blood infection, such as is the case for *P. falciparum*^7–10^. Clinically, a compound with radical cure efficacy is one that removes all parasites from the patient, including hypnozoites in the liver^11^.

Hypnozoites are refractory to all antimalarials except the 8-aminoquinolines, which were first identified over 70 years ago using low-throughput screening in avian malaria models^12^. Primaquine was the first 8-aminoquinoline widely-used for radical cure, however, efficacy is contingent on a large total dose administered in a 7-14 regimen, leading to adherence problems and infrequent use in malaria control programs of endemic countries^13^. Tafenoquine-chloroquine was developed from primaquine as an improved single-dose for radical cure^14^, but a recent clinical trial shows tafenoquine lacks efficacy when co-administered with the common antimalarial dihydroartemisinin-piperaquine, calling into question tafenoquine’s suitability in areas of high chloroquine resistance^15^. Furthermore, 8-aminoquinolines cannot be administered to pregnant women or glucose-6-phosphate dehydrogenase-deficient individuals and are ineffective in persons with specific cytochrome P450 genotypes^16^. For these reasons, the discovery and development of new chemical classes with radical cure activity are needed^17^.

Modern drug discovery typically relies on phenotypic screening and protein target identification^18^. For malaria, this approach ensures hits are acting on parasite targets and enables rational drug design, leading to several promising novel classes of antimalarials^19,20^. However, due to lower cost and higher feasibility, current high throughput screening for new antimalarials focuses almost entirely on blood or liver schizonts^21,22^. High throughput antimalarial screening with a target chemo-profile for killing hypnozoites has only recently been made possible with the introduction of cell-based phenotypic screening platforms featuring a monolayer of hepatocytes infected with sporozoites, a portion of which go on to form hypnozoites^23^. While the first hypnozonticidal hits from these platforms are just now being reported^24^, protein target identification approaches for hypnozonticidal drug discovery are in their infancy as the transcriptome of hypnozoites has only recently been reported and robust methods for genetic manipulation of *P. vivax* are still underdeveloped^25,26^.

To address the lack of radical cure drug leads and targets, we used our advanced *P. vivax* liver stage platform to first screen the Repurposing, Focused Rescue, and Accelerated Medchem (ReFRAME) library^7^. This library consists of approximately 12,000 developmental, approved, and discontinued drugs with the expectation that the repurposing of compounds with some optimization or regulatory success could expedite the decade-long path drugs typically progress through from discovery to licensure^27^. To accomplish this screen, we assembled an international collaboration with laboratories in malaria-endemic countries whereby vivax-malaria patient blood was collected and fed to mosquitoes to produce sporozoites for infecting primary human hepatocytes (PHH) in screening assays performed on-site. Interestingly, two structurally related compounds used for treating hypertension, hydralazine and cadralazine, were found effective at killing hypnozoites. Because these inhibitors have been shown to modulate DNA methylation^28,29^, we pursued and confirmed the existence of methyl-cytosine modifications in *P. vivax* sporozoite and liver stages. Having found in the ReFRAME screen a class of hits targeting an epigenetic pathway, we decided to confirm the importance of epigenetics in *P. vivax* hypnozoites and screened an additional commercial epigenetic inhibitor library using an improved version of our screening platform. Hypnozoites were found to be susceptible to several classes of epigenetic inhibitors, including six distinct histone deacetylase inhibitors and two inhibitors targeting histone methylation. To further assess the importance of histone acetylation in *P. vivax* liver stages, we tested inhibitors previously reported to be directly acting on *P. falciparum* acetyl-CoA synthetase, thereby modulating the pool of acetyl-CoA available for histone acetylation^30^. We found MMV019721 selectively kills *P. vivax* and *P. cynomolgi* hypnozoites, implicating acetyl-CoA synthetase is an additional hypnozonticidal drug target. This work demonstrates that in lieu of traditional molecular biology methods, our screening platforms identify multiple, druggable epigenetic pathways in hypnozoites and adds to the growing body of evidence that epigenetic features underpin biology in *P. vivax* and *P. cynomolgi* sporozoite and liver stages^25,31–33^.

## Results

### ReFRAME library screening cascade, hit identification, and confirmation

Chemical biology approaches have shown that hypnozoites become insensitive to most legacy antimalarials after 5 days in culture, indicating they must mature following hepatocyte infection^24,34^. Hypnozoite maturation was also noted in recent single-cell transcriptomic analyses of *P. vivax* liver stages, which demonstrate distinct population clusters of maturing and quiescent hypnozoites^10,25^. Importantly, discovery and development of hit compounds with radical cure activity *in vivo*, which includes elimination of hypnozoites in the liver of malaria patients^11^, requires screening against mature hypnozoites *in vitro*^35^. While our 8-day *P. vivax* liver stage platform, in which sporozoites are infected into primary human hepatocytes (PHH) and then allowed to mature for 5 days before being treated with test compound^36^, has been used for screening small libraries against mature hypnozoites^24^, the size of the ReFRAME library (12,823 compounds tested at 10 μM) presented a logistical challenge. We anticipated that dozens of *P. vivax* cases, each with a unique genetic background, would be needed to produce the sporozoites required to screen the 40 microtiter plates containing the library. To preclude the complex process of regular international shipments of infected mosquitoes, the *P. vivax* liver stage platform was successfully adapted and set up in research labs in two distinct malaria endemic areas, the Shoklo Malaria Research Unit (SMRU) in Thailand and the Institute Pasteur of Cambodia (IPC). The screening library was also divided between both sites to enable concurrent progress; ultimately 36 *P. vivax* cases from either site were needed to complete the primary screen over the course of 18 months (Figs. 1A, S1).

**Fig. 1.**
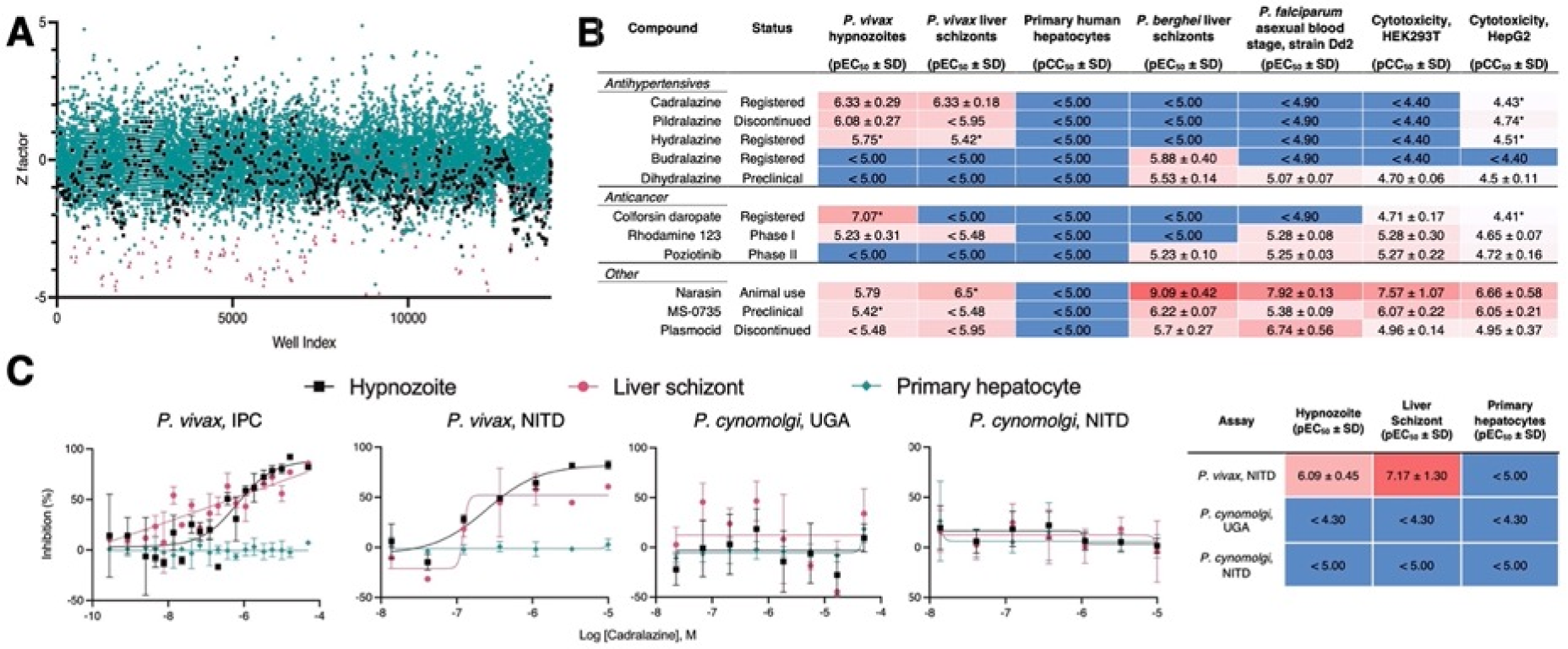
Hypnozonticidal hit detection and confirmation. (**A**) Index chart depicting the primary screen of the ReFRAME library against *P. vivax* hypnozoites in an 8-day assay. Hypnozoite counts were normalized by mean quantity per well for each plate (Z factor). Teal: library, black: DMSO, red: 1 μM monensin. (**B**) Primary screen hits were confirmed by dose-response in 8-day *P. vivax* liver stage assays and counterscreened against *P. berghei* liver schizonts, *P. falciparum* asexual blood stages (strain Dd2), HEK293T, and HepG2. Values represent pEC_50_ or pCC_50_ ± SD of all independent experiments (n=2-6) for which a pEC_50_ or pCC_50_ was obtained. (**C**) Dose-response curves for cadralazine against *P. vivax* and *P. cynomolgi* liver forms in 8-day assays at the IPC, UGA, and NITD. (B,C) Heat maps represent red as more potent and blue as inactive at highest dose tested. Asterisk (*) indicates only one independent experiment resulted in a calculated pEC_50_ or pCC_50_ (pEC_50_ is the inverse log of potency in M concentration, e.g. pEC_50_ 3 = 1 mM, pEC_50_ 6 = 1 μM, and pEC_50_ 9 = 1 nM). (C) All replicate wells were plotted together from all independent experiments (n=3 for *P. vivax* at IPC, n=1 for *P. vivax* at NITD, n=2 for *P. cynomolgi* at UGA, and n=4 for *P. cynomolgi* at NITD), bars represent SEM.

Some hits exhibited moderate selectivity and potency, with pEC_50_’s ranging from 5.42-7.07 (pEC_50_ is the inverse log of potency in M concentration, e.g. pEC_50_ 3 = 1 mM, pEC_50_ 6 = 1 μM, and pEC_50_ 9 = 1 nM) (Figs. 1B, S2). Colforsin daropate, rhodamine 123, and poziotinib are used to treat cancer and have known human targets, indicating that the targeted host pathways may be critical for hypnozoite persistence. As an example, poziotinib inhibits HER2, a tyrosine protein kinase associated with the downregulation of apoptosis and metastasis^37^. We recently reported that host apoptotic pathways are downregulated in *P. vivax-*infected hepatocytes^25^. Poziotinib could therefore act by upregulating apoptotic pathways in infected host cells. MS-0735, an analog of our previously reported hypnozonticidal hit, MMV018983^24^, is a ribonucleotide-reductase (RNR) inhibitor and used as an antiviral. The apparent need for nonreplicating hypnozoites to produce deoxyribonucleosides for DNA synthesis is peculiar. However, it has been reported that RNR is also critical for DNA damage repair^38^, is important for maintaining cancel cell dormancy^39^, and is expressed in *P. vivax* liver schizonts and hypnozoites^25^. We also rediscovered previously-reported hypnozonticidal compounds included in the library, including the ionophore narasin^24^ and the 8-aminoquinoline plasmocid^40^.

From our analysis of primary screen activity, we noted several hydrazinophthalazine vasodilators were potentially active (Fig. S1C). We selected 72 compounds for confirmation of activity against hypnozoites in a dose-response format, including 10 hydrazinophthalazine analogs. These compounds were counter-screened for additional antimalarial activity against *P. falciparum* blood stages and *P. berghei* liver schizonts. They were also tested for cytotoxicity against HEK293T and HepG2 human cell lines (Figs. 1B-C, S2). We confirmed that three hydrazinophthalazines analogs--cadralazine, pildralazine, and hydralazine--were active against mature hypnozoites, with cadralazine displaying the best combination of potency (pEC_50_ = 6.33 ± 0.33), maximal inhibition near 100%, and selectivity over PHH (> 21 fold), HEK293T (> 85 fold), and HepG2 (> 79 fold) cells (Figs. 1, B-C, S2A). Hydralazine, which was FDA-approved in 1953, is currently one of the world’s most-prescribed antihypertensives, and on the WHO list of essential medicines^41^. Cadralazine, which was developed in the 1980’s as an improvement over hydralazine, was abandoned due to side effects and only licensed in Italy and Japan^42^. Hydrazinophthalazines have been shown to inhibit human DNA methyltransferases (DNMT)^28,29^ and hydralazine has also been recently used to study potential DNA methylation patterns in the *P. falciparum* asexual blood stages^43^. Similar to our previous report^43^, these hydrazinophthalazines were inactive when tested against *P. berghei* liver schizonts and *P. falciparum* asexual blood stages, suggesting that hypnozoite quiescence may be biologically distinct from developing schizonts^24^. While hydrazinophthalazines may act on infected hepatocytes and not directly on the parasite, their distinct selectivity suggests that their effect is likely on a host or parasite pathways and not simply due to cytotoxicity in the host cell. Hydralazine and cadralazine were not identified as potential hits in any of the 112 bioassay screens of the ReFRAME published to date^44^, suggesting these compounds specifically target *P. vivax* liver stages and not promiscuously active compounds.

Methods for the robust culture of *P. vivax* hypnozoites were only recently reported, leading to several new reports on hypnozoite biology and radical cure drug discovery^7,45^. Consequentially, some hypnozoite-specific discoveries appear to be platform-specific^10,25^. Select hits were shared with the Novartis Institute for Tropical Diseases (NITD), where the activity and potency of cadralazine (pEC_50_ = 6.09 ± 0.45), hydralazine (pEC_50_ = 6.20), and poziotinib (pEC_50_ = 6.17) were independently confirmed in a similar 8-day *P. vivax* screening platform using a *P. vivax* case from southern Thailand. (Figs. 1C, S3A). Independent confirmation of these hits indicates their activities are not merely platform-specific and are, rather, more broadly descriptive of hypnozoite chemo-sensitivity.

Following our screening and hit confirmation, we investigated the potency, *in vivo* stability, and tolerability profile of our confirmed hits and chose cadralazine and hydralazine for repurposing as radical cure antimalarials. Currently, the gold-standard model for preclinical assessment of *in vivo* anti-relapse efficacy is rhesus macaques infected with *Plasmodium cynomolgi*, a zoonotic, relapsing species closely related to *P. vivax*^46^. Because we found cadralazine substantially more potent against hypnozoites than hydralazine, it was selected for a rhesus macaque pharmacokinetic study in which plasma levels were measured over 24 h following an oral dose of 1 mg/kg, which was calculated to be well-tolerated, and 30 mg/kg, which was calculated to likely cause drug-induced hypotension^47–49^. The 30 mg/kg dose resulted in maximum plasma concentration of 13.7 μg/mL (or 48.2 μM) and half-life of 2.19 ± 0.24 h, which was sufficient to cover the *in vitro* EC_90_ for several hours without noticeable side effects. (Fig. S4). As another prerequisite for *in vivo* validation, we next sought to confirm and measure the potency of cadralazine and other ReFRAME hits against *P. cynomolgi* B strain hypnozoites *in vitro* using an 8-day assay featuring primary simian hepatocytes (PSH) at NITD. While poziotinib was active against *P. cynomolgi* hypnozoites when tested two of three different PSH donor lots (pEC_50_ = 5.67 and 5.95), hydralazine and cadralazine were found inactive when tested in all three different PSH donor lots (Figs. 1C, S3B). This negative result was later confirmed in an 8-day, simianized version of the platform at the University of Georgia (UGA) using the *P. cynomolgi* Rossan strain infected into two different PSH lots (Fig. 1C). Altogether these data highlight potential difference between *P. vivax* and *P. cynomolgi* and challenge the gold-standard model for preclinical assessment of *in vivo* anti-relapse efficacy is rhesus macaques.

### Synergy between cadralazine and 5-azacytidine

As molecular tools to validate drug target in *P. vivax* are limited, we further confirmed the possible mechanism of action of hydrazinophthalazines using drug combination studies to assess synergy, additivity or antagonsim^30^. We used 5-azacytidine, a known DNA methyltransferase inhibitor^50^, to investigate its effects on cadralazine treatment. When tested alone in dose-response from 50 μM, 5-azacytidine had no effect on hypnozoites. However, when added to cadralazine in fixed ratio combinations ranging from 8:1 to 1:8, 5-azacytdine increased the potency of cadralazine by ∼2 fold across several combinations in two independent experiments (Figs. 2, S5). The most potent effect was detected using a 2:1 fixed ratio of cadralazine:5-azacytidine, resulting to an equivalent EC_50_ decrease from 470 nM to 216 nM.

**Fig. 2.**
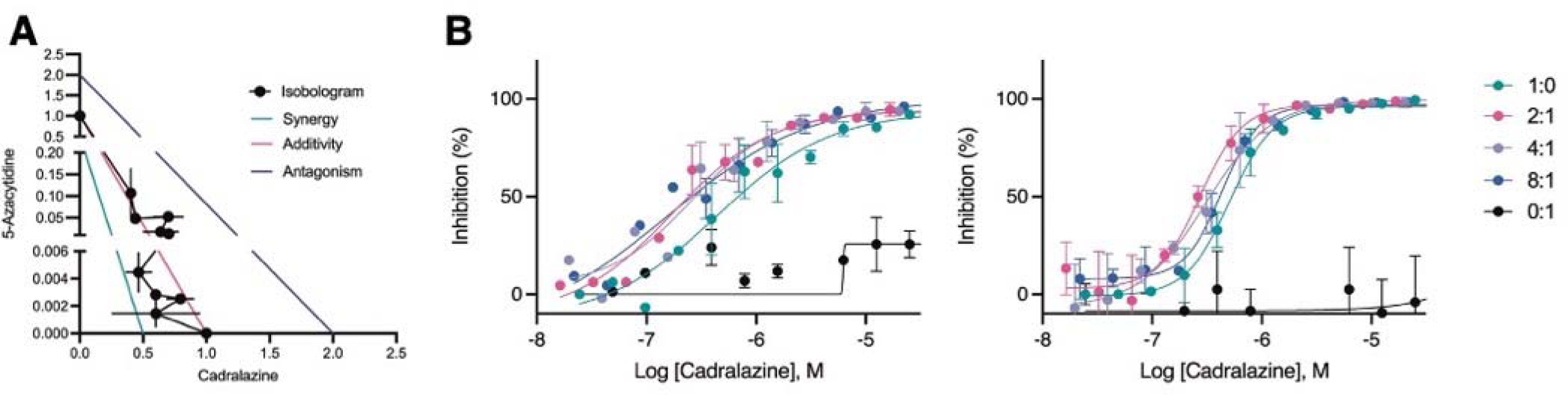
Synergistic effect of cadralazine and 5-azacytidine in *P. vivax* liver stage assays. (**A**) Isobologram of cadralazine and 5-azacytidine activity against hypnozoites in fixed ratios of 1:0, 8:1, 6:1, 4:1, 2:1, 1:1, 1:2, 1:4, 1:6, 1:8, and 0:1, bars represent SD of FICs from two independent experiments. (**B**) Dose-response curves for cadralazine at the most synergistic fixed ratios (2:1, 4:1, and 8:1) against hypnozoites. Cadralazine alone is represented as 1:0, 5-azacytidine alone is represented as 0:1 and plotted on the cadralazine chart for comparison. Left and right charts represent two independent experiments, bars represent replicate wells at each dose.

### Immunofluorescent detection of DNA methylation in *P. vivax* and *P. cynomolgi* liver stages

To further confirmed that cadralazine interacts with *P. vivax* target(s), we aimed to detect and quantify DNA methylation in the *P. vivax* and *P. cynomolgi* genomes. Previous studies had identified the presence of low level 5-methylcytosine (5mC), 5-hydroxmethylcystosine (5hmC), and 5hmC-like marks throughout the genome^43,51,52^. We first conducted an immunofluorescence staining assay using commercially available anti-5mC and anti-5hmC monoclonal antibodies to identify evidence of DNA methylation in *P. vivax* liver stages at 6 days post-infection. We found clear evidence of 5mC, but not 5hmC, in both schizonts and hypnozoites, morphologically consistent with the presence of 5mC in the parasite’s nucleus^7^ (Figs. 3A, S6-8). To segregate signals coming from the host hepatic nuclei, we used automated high content imaging analysis on hundreds of individual *P. vivax* liver stage parasites as an unbiased approach for quantifying 5mC signal within parasites. Image masks were generated to quantify the area of 5mC or 5hmC stain within each parasite (Fig. S9). The values were then plotted as stain area per hypnozoite or per schizont (Fig. 4B). While some evidence of 5hmC-positive forms did appear from this analysis, the net 5hmC area per parasite was found significantly lower when compared to 5mC signals (Kurskal-Wallis tests, for hypnozoites *H*(7) = 194.3, *p* <.0001, for schizonts *H*(7) = 88.66, *p* <.0001). Similar results on the ratio of 5hmC to 5mC were also recently reported in *P. falciparum* blood stages^53^, confirming that 5mC marks are the predominate DNA methylation marks in both species.

**Fig. 3.**
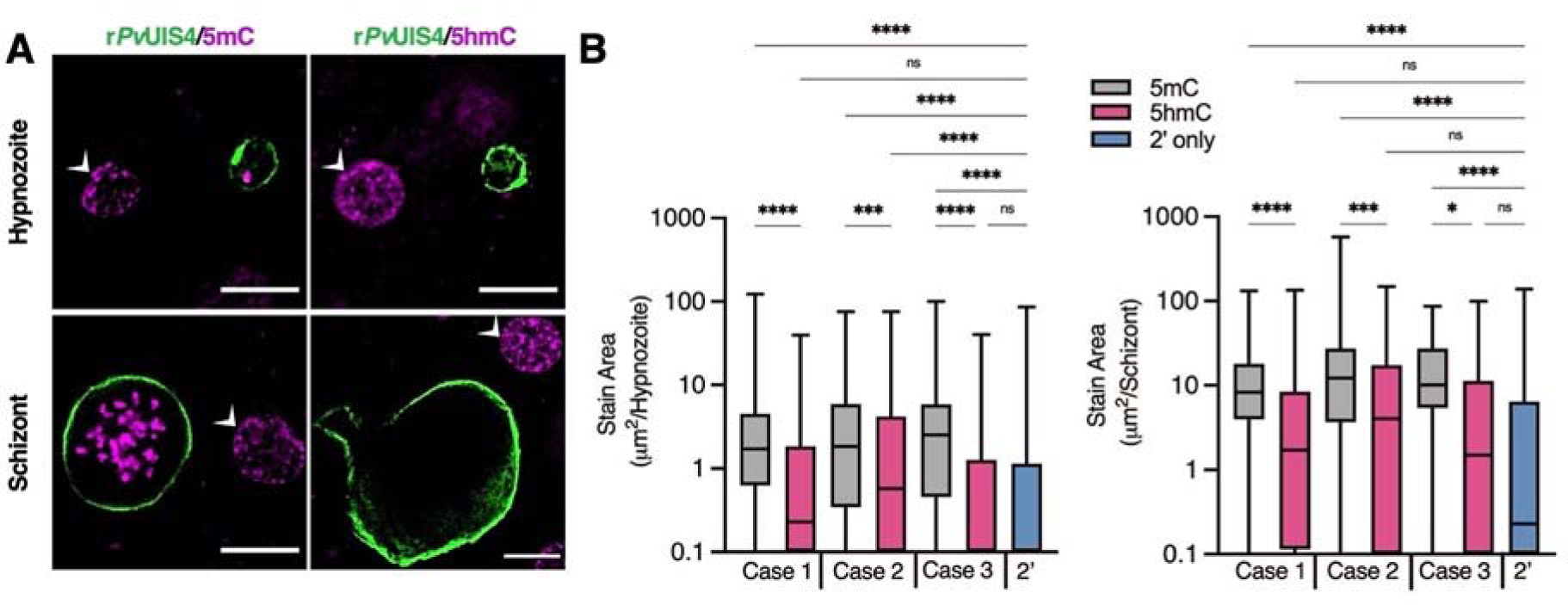
Cytosine modifications in *P. vivax* liver forms. (**A**) Immunofluorescent imaging of a 5mC-positive (left) or 5hmC-negative (right) *P. vivax* hypnozoite (top) and schizont (bottom) at day 6 post-infection. White arrows indicate hepatocyte nuclei positive for 5mC or 5hmC. Bars represent 10 µm. (**B**) High-content quantification of 5mC or 5hmC stain area within hypnozoites or schizonts from sporozoites generated from three different *P. vivax* cases. Significance determined using Kruskal-Wallis tests, for hypnozoites *H*(7) = 194.3, *p* <.0001, for schizonts *H*(7) = 88.66, *p* <.0001, with Dunn’s multiple comparisons, *\*p* <.05, \*\*\**p* <.001, \*\*\*\**p* <.0001, ns = not significant. Line, box and whiskers represent median, upper and lower quartiles, and minimum-to-maximum values, respectively, of all hypnozoites (177 ≤ n ≤ 257) or all schizonts (30 ≤ n ≤ 142) in culture for each case, 2’ indicates a secondary stain only control.

**Fig. 4.**
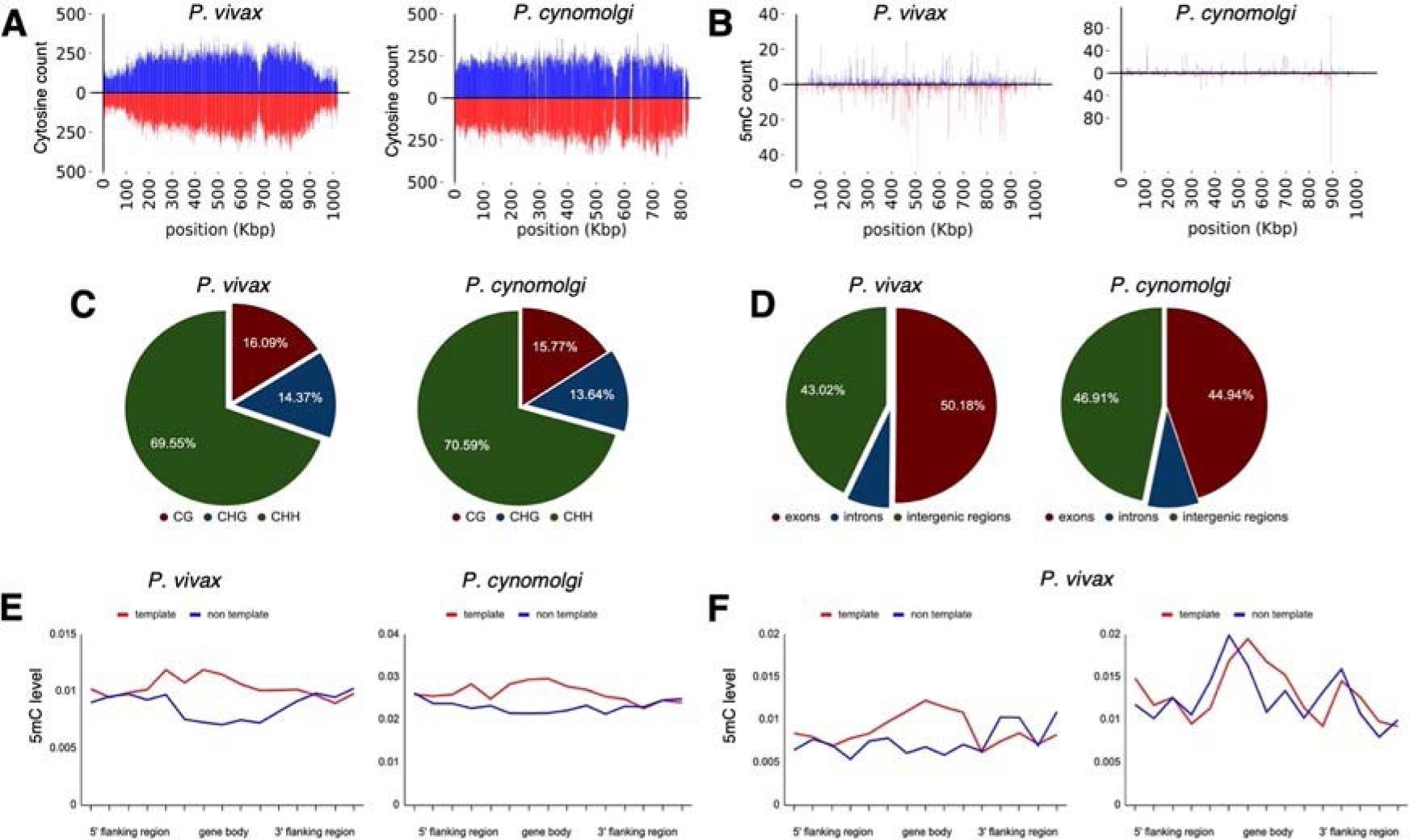
Density of cytosine and methylated cytosine (5mC) in sporozoites. (**A**) CG content of chromosome 1 for *P. vivax* and *P. cynomolgi*. The total number of cytosines was quantified on each strand using 1 kbp long non-overlapping windows. (**B**) The total number of methylated cytosines was quantified on each strand using 1 kbp long non-overlapping windows. (**C**) The number of 5mC present in all possible contexts (CG, CHG, and CHH) quantified throughout the genome of *P. vivax* and *P. cynomolgi.* (**D**) Repartitioned 5mC quantity within different compartments of the genome in *P. vivax* and *P. cynomolgi.* (**E**) Strand-specificity of 5mC for all genes in *P. vivax* and *P. cynomolgi*. Flanking regions and gene bodies were divided into five bins and the methylation level of each bin was averaged among all genes. Red: template strand, blue: non-template strand. (**F**) The previously reported mRNA abundance of *P. vivax* sporozoites was retrieved^22^ and genes ranked. The 5mC levels in 5’ flanking regions, gene bodies, and 3’ flanking regions were placed into five bins and are shown for highly expressed (90th percentile, left) and weakly expressed (10th percentile, right) genes. Red: template strand, blue: non template strand.

Given the different susceptibility of *P. cynomolgi* hypnozoites to hydrazinophthalazines as compared to *P. vivax*, we performed automated high content analysis of 5mC-and 5hmC-stained *P. cynomolgi* M/B-strain liver schizonts and hypnozoites at 8 and 12 days post-infection. Like *P. vivax*, we found both *P. cynomolgi* liver schizonts and hypnozoites are positive for 5mC, but not 5hmC. However, the 5mC stain morphology and intensity were relatively lower in *P. cynomolgi* hypnozoites versus *P. vivax* hypnozoites, suggesting potential divergence of DNA methylation pathways in these two species (Fig. S10).

### Detection of cytosine modifications in *P. vivax* and *P. cynomolgi* sporozoites using liquid chromatography-tandem mass spectrometry and bisulfite sequencing

We next sought to confirm the presence of cytosine methylation in the *P. vivax* and *P. cynomolgi* genomes using mass spectrometry and bisulfite sequencing. We initially assessed that without an available single cell sequencing approach, sequencing coverage of the parasite’s genome would be overwhelmed by the genomic material from the host cell as well as neighboring uninfected hepatocytes^25^. We therefore collected sufficient genomic material from *P. vivax* and *P. cynomolgi* sporozoites to analyze the nucleoside mixture arising from the enzymatic digestion of genomic DNA by liquid chromatography-tandem mass spectrometry as well as for detection of DNMT activity using a commercial *in vitro* DNA methylation assay^43^. While we detected 5mC and DNMT activity in *Plasmodium-*enriched samples with these approaches, possible contamination by the mosquito’s microbiota could not be excluded (Fig. S15A-B). We next analyzed DNA methylation loci at single-nucleotide resolution using bisulfite sequencing of 3×10^7^ *P. vivax* sporozoites, generated from three different cases, as well as 3×10^7^ *P. cynomolgi* sporozoites (Fig. 4, A-B). A total of 161 and 147 million high-quality reads were sequenced for *P. vivax* and *P. cynomolgi* samples, respectively (Fig. S11C). The average 5mC level detected across all cytosines was 0.49% and 0.39% for *P. vivax* and *P. cynomolgi*, respectively. These percentages are comparable to the 0.58% methylation level detected in *P. falciparum* blood stages^43^, but likely underestimate methylated loci considering the coverage we achieved (see methods).

We then monitored the distribution of detected 5mC along the *P. vivax* and *P. cynomolgi* chromosomes (Figs. S12-13) and observed a stable methylation level throughout the genomes, including in telomeric and sub-telomeric regions. We further examined the context of genome-wide methylations and, similar to what we previously observed in *P. falciparum*^43^, methylation was detected as asymmetrical, with CHH (where H can be any nucleotide but G) at 69.5% and 70.5%, CG at 16% and 15.7%, and CHG at 14.3% and 13.64%, for *P. vivax* and *P. cynomolgi*, respectively (Fig. 4C). We then measured the proportion of 5mC in the various compartments of gene bodies (exons, the introns, promoters, and terminators) as well as strand-specificity (Fig. 4, D-E). We observed a slightly increased distribution of 5mC in promoters and exons compared to the intronic region, as well as in the template versus non-template strand, in *P. vivax* and *P. cynomolgi*. These results were consistent with previous data obtained in *P. falciparum* and in plants^43,52^. Such a strand specificity of DNA methylation patterns can affect the affinity of the RNA polymerase II and impact transcription, thus we compared methylation levels to previously-report transcriptomic data from *P. vivax* sporozoites^32^. The 5mC levels in 5’ flanking regions, gene bodies, and 3’flanking regions were placed into five bins and compared to mRNA abundance, revealing an inverse relationship between methylation and mRNA abundance in the proximal promoter regions and the beginning of the gene bodies, with highly-expressed genes appearing hypomethylated and weakly-expressed genes hypermethylated (Fig. 4F). These results suggest that methylation level in proximal promoter regions as well as in the first exon of the genes may affect, at least partially, gene expression in malaria parasites. While these data will need to be further validated and linked to hypnozoite formation at a single-cell level, we have determined that 5mC is present at a low level in *P. vivax* and *P. cynomolgi* sporozoites and could control liver stage development and hypnozoite quiescence.

### Assay improvements and epigenetic inhibitor library screen

The success of the original screening platform protocol and secondary confirmation of several of our initial hits provided us an invaluable opportunity to develop an improved radical cure screening assay. The current iterations of our screening platform rely on high-content analysis of parasitophorous vacuole staining of the forms that persist up to the assay endpoint^7,54^. During the course of the ReFRAME primary screen we found the day 8 endpoint was sufficient for some hit compounds to act. However, other compounds like the 8-aminoquinolines exhibit a ‘delayed death’ phenotype, which leads to a false-negative result^24^. We therefore extended the assay by 4 days to allow attenuated forms to be cleared from the culture^24^. Also, as our screening assays were performed with multiple lots of PHH and PSH, we detected some lot-specific results, including small differences in activity of poziotinib and our monensin control (Figs. 1B, S3, S14A), possibly due to compound instability in the presence of hepatic metabolism. We therefore tested the metabolism inhibitor 1-aminobenzotriazole (1-ABT) in culture media to minimize the effect of lot-specific hepatic metabolism^55^. We used a cytochrome P450 functional assay specific to CYP3A4 and determined that 100 μM of 1-ABT was sufficient to completely reduce CYP3A4 activity in both basal and rifampicin-induced PHH (Fig. S15A-B). This effect was further confirmed and quantified by mass spectrometry after 1 h of treatment at 100 μM 1-ABT. We not only detected a 75% decrease in CYP3A4 activity, but also a more than 60% reduction of CYP2B6 and CYP2E1 activity along with lesser effects on CYP2C9, CYP1A2, and CYP2D6 (Fig. S15C). These changes were incorporated into our original 8-day protocol to design an improved 12-day assay^36^ that we then validated by re-testing 12 ReFRAME hits. The modified assay did not drastically affect the potency of most hits (Fig. S16A), but confirmed hypnozonticidal activity for poziotinib (pEC_50_ = 6.05), which had only been previously confirmed in *P. vivax* and *P. cynomolgi* assays performed at NITD only (Figs. 1B, S3, S16B). This assay was then use in all followup experiments.

To further confirm the importance of epigenetics in hypnozoites biology,^31^ we obtained a commercially-available library containing 773 compounds targeting various inhibitors of epigenetic enzymes or pathways. These compounds were tested at 10 μM against *P. vivax* liver stages at both SMRU and IPC sites (Fig. S17A). We confirmed our initial hits in dose-response assays resulting in selective hypnozonticidal potency for 11 compounds targeting five different epigenetic mechanisms (Tab. 1, Fig. S17B). This includes the histone deacetylase inhibitors panobinostat (pEC_50_ = 6.98 ± 0.18), AR42 (pEC_50_ = 6.11 ± 0.24), abexinostat (pEC_50_ = 5.48 ± 0.00), givinostat (pEC_50_ = 5.35 ± 0.45), practinostat (pEC_50_ = 5.32 ± 0.13), and raddeanin A (pEC_50_ = 5.95 ± 0.00). Histone methyltransferase inhibitor hits included MI2 (pEC_50_ = 5.48 ± 0.00), a compound that targets the interaction between menin (a global regulator of gene expression), and MLL (a DNA-binding protein that methylates histone H3 lysine 4^56^), and cyproheptadine (pEC_50_ = 5.24 ± 0.34), which targets the SET-domain-containing lysine methyltransferase^57^. These results corroborate our hypothesis that epigenetic pathways regulate hypnozoites^31,32^. Other hits, including 666-15 (pEC_50_ = 5.88 ± 0.12), an inhibitor of the transcription factor cAMP response element-binding protein^58^, and cerdulatinib (pEC_50_ = 5.33 ± 0.20), a kinase inhibitor, suggest that signaling pathways may also be important for quiescence^59^.

**Table 1.**
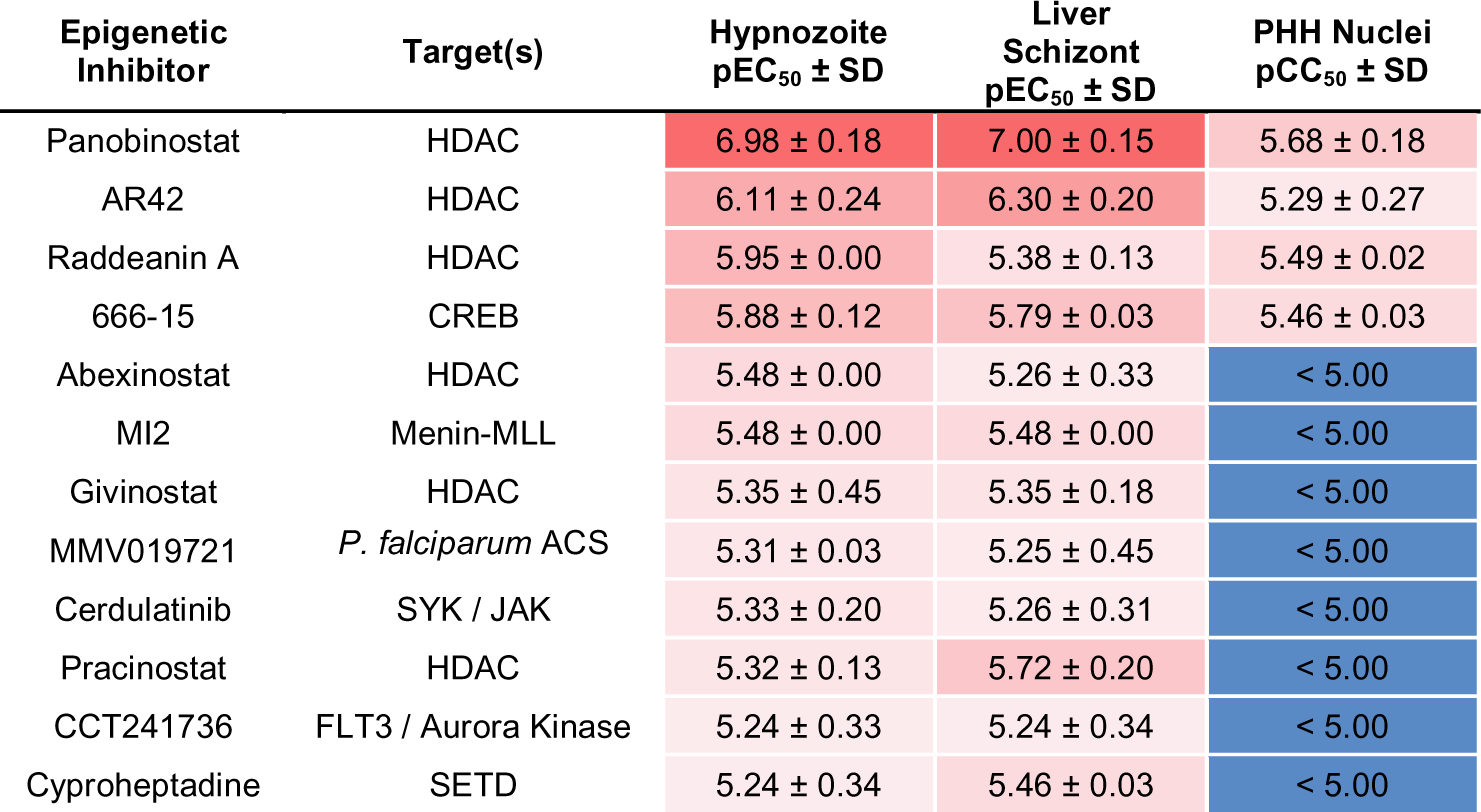
Additional epigenetic inhibitors with activity against *P. vivax* liver stages. HDAC: histone deacetylase. CREB: cAMP response element-binding protein. FLT3: fms-like tyrosine kinase 3. *P. falciparum* ACS: *P. falciparum* acetyl CoA synthetase. SYK: spleen tyrosine kinase. JAK: Janus kinase. SETD: SET domain containing histone lysine methyltransferase. Mean and standard deviation are from two or more independent experiments.

| Epigenetic Inhibitor | Target(s) | Hypnozoite pEC<sub>50</sub> ± SD | Liver Schizont pEC<sub>50</sub> ± SD | PHH Nuclei pCC<sub>50</sub> ± SD |
| --- | --- | --- | --- | --- |
| Panobinostat | HDAC | 6.98 ± 0.18 | 7.00 ± 0.15 | 5.68 ± 0.18 |
| AR42 | HDAC | 6.11 ± 0.24 | 6.30 ± 0.20 | 5.29 ± 0.27 |
| Raddeanin A | HDAC | 5.95 ± 0.00 | 5.38 ± 0.13 | 5.49 ± 0.02 |
| 666-15 | CREB | 5.88 ± 0.12 | 5.79 ± 0.03 | 5.46 ± 0.03 |
| Abexinostat | HDAC | 5.48 ± 0.00 | 5.26 ± 0.33 | < 5.00 |
| MI2 | Menin-MLL | 5.48 ± 0.00 | 5.48 ± 0.00 | < 5.00 |
| Givinostat | HDAC | 5.35 ± 0.45 | 5.35 ± 0.18 | < 5.00 |
| MMV019721 | *P. falciparum* ACS | 5.31 ± 0.03 | 5.25 ± 0.45 | < 5.00 |
| Cerdulatinib | SYK / JAK | 5.33 ± 0.20 | 5.26 ± 0.31 | < 5.00 |
| Pracinostat | HDAC | 5.32 ± 0.13 | 5.72 ± 0.20 | < 5.00 |
| CCT241736 | FLT3 / Aurora Kinase | 5.24 ± 0.33 | 5.24 ± 0.34 | < 5.00 |
| Cyproheptadine | SETD | 5.24 ± 0.34 | 5.46 ± 0.03 | < 5.00 |

Having identified several histone deacetylase inhibitors as directly or indirectly active on hypnozoites, we screened compounds previously reported as inhibitors of *P. falciparum* acetyl-CoA synthetase (ACS), with downstream effects on histone acetylation^30^. We found that one compound, MMV019721, was selectively active on mature *P. vivax* hypnozoites (Tab. 1). Given the evidence MMV019721 is directly targeting *P. falciparum* ACS, this result suggests ACS also is a hypnozonticidal drug target. While the molecular techniques needed to confirm the direct interaction of MMV019721 and ACS in *P. vivax* are currently underdeveloped, our data supplement recent reports describing epigenetics as important regulators in *P. vivax and P. cynomolgi* at different stage of the parasite life cycle^25,32,33^.

## Discussion

Herein we demonstrate several significant advances that progress radical cure antimalarial drug discovery and development, including the first report of screening a medium-sized (>10,000) compound library against mature hypnozoites as well as detection of novel hits with mechanisms unrelated to that of 8-aminoquinolines. Identification of these hits was made possible following the establishment of a complex logistical operation in which the sporozoites used for screening were produced by feeding *P. vivax*-infected blood from malaria patient isolates to mosquito colonies at malaria research institutes in two countries in Southeast Asia. Our international collaboration overcame several logistical hurdles to obtain positive Z’ scores for most screening plates. Hits were also confirmed via dose-response, indicating that expanded screening directed against *P. vivax* liver stages is likely to produce more hypnozoite-specific hits (Figs. 1, 5, S1A-B, S17).

The only class of FDA-approved compounds for radical cure, the 8-aminoquinolines, were not discovered from *in vitro* drug screening. Instead, they were discovered using animal models, including the *P. cynomolgi-*infected rhesus macaque system^12^. The 8-aminoquionlines function through generation of reactive oxygen species affecting both the host and parasite, and lack a distinct parasite target^60–63^. As such, this work represents one of the first applications of a radical cure development pipeline to begin with *in vitro* screening against *P. vivax* hypnozoites and end with attempted confirmation using *P. cynomolgi* radical cure models. While our screen generated positive results against *P. vivax,* we found mixed results against *P. cynomolgi* hypnozoites *in vitro* (Figs. 1C, S3B). While further studies will be needed to confirm that target(s) of our hits are parasite-or host-directed and may influence parasite survival, our data show there is sufficient diversity in gene expression, structural biology, or mechanisms of hepatic quiescence between *P. cynomolgi* and *P. vivax* hypnozoites that some newly identified hits may be species-specific. While this result could also be attributed to differential metabolism in human and monkey hepatocytes^64^, the rhesus macaque radical cure model is currently considered as an important prerequisite for continued drug development, including efficacy testing in controlled human infections. The role of this model in the radical cure drug development cascade may need to be reevaluated as some compounds identified as promising for the radical cure of *P. vivax* may be abandoned too quickly due to the lack of activity against *P. cynomolgi.* This result highlights the need for further development and validation of *P. vivax*-specific animal models^65^. As a whole this report adds to the broader discussion surrounding the successes and challenges of drug repurposing^66^. While direct repositioning of a known drug as a safe treatment for a new indication is the ideal outcome, it can serve as advanced starting points for further optimization and has still the potential for reducing the time and cost involved in developing an efficacious therapy.

In addition to the identification of promising new hits and direction, we also confirmed that epigenetic control of pathogenic dormancy via DNA methylation is a pathway that could be potentially targeted by future antimalarials. This pathway has already been described for several disease agents capable of dormancy, including cancer cells^67^ and tuberculosis^68^. DNA methylation has also been validated as controlling critical processes in plants, which share evolutionary traits with *Plasmodium*^69^. DNA methylation in the genus *Plasmodium* was first described in *P. falciparum* blood stages^43^, and has been associated with gene expression, transcriptional elongation and parasite growth^51,52,70^. Previous experiment have shown that hydralazine can directly inhibits DNA methylation in nuclear extracts of blood stage parasites but also inhibit a recombinant functional fragment of the *P. falciparum* DNMT^43^. We pursued several biomolecular approaches to confirm that cadralazine may also interact with *P. vivax* DNMT in liver stage parasites. Due to technical limitations, we used a two-drug combination study in which the known DNMT inhibitor 5-azacytidine potentiated cadralazine against *P. vivax* hypnozoites (Figs. 2, S5). While we continue to develop new protocols and confirm the direct interaction of cadralazine with *P. vivax*, we successfully confirmed 5mC marks in *P. vivax* and *P. cynomolgi* liver stage parasites using both immunofluorescence and whole genome bisulfite sequencing assays (Figs. 3-4).

The current model of hypnozoite quiescence suggests RNA binding proteins (RBPs) drive hypnozoite formation by preventing translation of target mRNAs associated with schizogony^33^. In this model, histone acetylation results in euchromatin at the loci of RBPs, resulting in their expression and ongoing quiescence. Hypothetically, HDAC inhibitors would favor quiescence while a treatment that decreases histone acetylation would favor schizongony. This model somewhat contrasts with our present finding that HDAC inhibitors and the ACS inhibitor MMV019721 successfully kill hypnozoites *in vitro* (Tab. 1). It is, however, likely that the identified RBPs are part of broader gene networks which, when perturbed by sudden modulation of epigenetic feature such as DNA methylation and histone acetylation, results in a lethal level of dysregulation. While we still need to develop *P. vivax* transgenic lines to successful study hypnozoite biology and further validate potential drug targets^71,72^, the chemical probes that we described in this report could be used in combination with single-cell technology to more precisely perturb hypnozoites and refine our understanding of epigenetic pathways regulating hypnozoite formation and survival.

## Acknowledgments

We thank the malaria patients of Thailand and Cambodia for participation in this study. We thank the Sporocore at UGA for generating *P. berghei*-infected mosquitoes. We are grateful to Calibr’s Compound Management and High Throughput Screening Groups for their assistance with this project. HCI data from drug studies was produced by the Biomedical Microscopy Core at UGA, supported by the Georgia Research Alliance. SMRU is part of the Mahidol Oxford Research Unit, supported by the Wellcome Trust of Great Britain (#220211). Material has been reviewed by the Walter Reed Army Institute of Research. There is no objection to its presentation and/or publication. The opinions or assertions contained herein are the private views of the author, and are not to be construed as official, or as reflecting true views of the Department of the Army or the Department of Defense. This publication includes data generated at the University of California, San Diego IGM Genomics Center utilizing an Illumina NovaSeq 6000 that was purchased with funding from a National Institutes of Health SIG grant (#S10 OD026929).

## Funding

Funding support was provided by the Bill & Melinda Gates Foundation (#OPP1107194 to Calibr, INV-031788 to CJJ, and #OPP1023601 to DEK), Medicines for Malaria Venture (RD/17/0042 and RD/15/0022 to BW and AV and RD/15/0022 to SPM and DEK), the National Institutes of Allergy and Infectious Diseases of the National Institutes of Health (#HHSN272201200031C to MRG and #1R01 AI136511 to KGLR) and the University of California, Riverside (#NIFA-Hatch-225935 to KGLR)

## Author contributions

Conceptualization: SPM, MAB, AV, AKC, CWM, BW, KGLR, DEK; Methodology: SPM, MAB, AV, CAC, VP-I, CR, CJJ, FN, BW, KGLR; Validation: SPM, MAB, ELF, KGLR; Formal Analysis: SPM, MAB, AAR, KGLR; Investigation: SPM, MAB, AV, ELF, MG, YA-K, ATC, EC, CAC, KC, AH-C, SBJ, TL, SL, JM, TM, AO, VP-I, KP, JPé, JPr, CR, SSe, SSu, YW, JY, KGLR; Resources: SPM, MAB, AV, CA, MA, MC-M, AKC, WTC, CAC, MRG, HJ, CR, SSS, CLS, JS, RU, PW, JP, CJJ, FN, BW; Data Curation: SPM, MAB, ELF, ATC, AH-C, KGLR; Visualization: SPM, MAB, JM, CR, KGLR; Funding acquisition: SPM, AV, MRG, CJJ, FN, BW, KGLR, DEK; Project administration: SPM, AV, FN, BW, DEK; Supervision: SPM, AV, ELF, BC, AKC, MG, SAM, JP, CJJ, FN, BW, KGLR, DEK; Writing – original draft: SPM, MAB, KGLR; Writing – review & editing: SPM, MAB, AV, ELF, CAC, KC, MRG, VP-I, CR, CJJ, FN, KGLR, DEK.

## Competing interests

TM and KC are employees of BioIVT. AH-C, ELF, and SAM are employees of the Novartis Institute for Tropical Disease, BC is an employee of MMV. All other authors have no competing interests.

## Data and materials availability

For the purpose of Open Access, the authors have applied a CC BY public copyright license to any Author Accept Manuscript version arising from this submission. All bisulfite sequencing data generated in this study can be found in the Sequence Read Archive (SRA) at the NCBI National Library of Medicine (https://www.ncbi.nlm.nih.gov/sra) under the BioProject code PRJNA925570.

## Supplementary Materials

### Materials and Methods

#### ReFRAME library description and plating

The ReFRAME library was curated by assembling a list of developmental and FDA-approved chemistry from three databases (GVK Excelra GoStar, Clarivate Integrity, and Citeline Pharmaprojects). The original library consisted of 36 384-well plates (ReF01-ReF36, Fig. S1A) containing 11,871 test compounds^27^. While the original library was being screened, an additional set of four 384-well plates (ReF38-ReF41, Fig. S1A) were added to the library, totaling 12,823 test compounds^44^. Source plates were made from the master library at Calibr at Scripps Research such that 3-5 μL of 10 mM solution was added to each well of a sterile, conical-bottom 384-well plate (Greiner Bio-One cat 784261). Most compounds were diluted in DMSO, however, a subset was diluted in water due to limited DMSO solubility. Plates were sealed and shipped on dry ice to SMRU and IPC and stored at −20 °C prior to use. Column 24 of each plate was filled with 5 μL DMSO to serve as negative control wells. Control compounds included 1 mM monensin (positive control for hypnozoite and schizont activity), 1 mM the phosphatidylinositol 4-kinase inhibitor (PI4Ki) KDU691 or MMV390048 (positive control for schizont activity), 1 mM atovaquone (negative control for radical cure activity) and 10 mM tafenoquine (clinically relevant control for hypnozoite activity)^7,24^.

#### Ethical Approval for Human Subjects and Animal Use

The Thai human subjects protocols for this study were approved by the Institutional Ethics Committee of the Thai Ministry of Public Health and the Oxford Tropical Medicine Ethical Committee (TMEC 14-016 and OxTREC 40-14). The Cambodian human subjects protocols for this study were approved by the Cambodian National Ethics Committee for Health Research (100NECHR, 104NHECR, 111NECHR, 113NHECR and 237NHECR). Protocols conformed to the Helsinki Declaration on Ethical Principles for Medical Research Involving Human Subjects^73^ and informed written consent was obtained for all volunteers or legal guardians. *P. cynomolgi* sporozoites were generated at Emory National Primate Research Center (ENPRC) using procedures approved by the Emory University Institutional Animal Care and Use Committee (PROTO201900110), as well as at UGA using procedures approved by UGA’s Institutional Animal Care and Use Committee (A2020 03-002-Y3-A15). *P. cynomolgi* sporozoites were also produced at the Armed Forces Research Institute of Medical Science under an IACUC-approved animal use protocol in an AAALAC International-accredited facility with a Public Health Services Animal Welfare Assurance and in compliance with the Animal Welfare Act and other federal statutes and regulations relating to laboratory animals (22-10). *P. berghei* sporozoites were generated by the Sporocore at UGA using procedures approved by UGA’s Institutional Animal Care and Use Committee (A2016 06–010–Y1–A0 and A2020 01-013-Y2-A3). Pharmacokinetic studies were conducted at WuXi AppTec Co., Ltd., in accordance with the WuXi IACUC standard animal procedures along with the IACUC guidelines that are in compliance with the Animal Welfare Act^74^.

#### ReFRAME primary screen against *P. vivax* liver stages

The complete, step-by-step protocol for the *P. vivax* liver stage assay is published^36^. In summary, two days after assay plates (Greiner Bio-One cat 781956) were seeded with PHH, sporozoites were dissected from mosquito salivary glands and allowed to infect cultures. The ReFRAME library was screened using the original, 8-day radical cure assay, in which developing liver schizonts and mature, PI4Ki-insensitive hypnozoites were treated on days 5-7 post-infection^7,24^. On treatment days, a pintool was used to transfer 40 nL of compounds from the source plates to the assay plates. A single PHH lot, UBV, was first used for screening, however, once all available cryovials were used, screening was completed with lot BGW (Fig. S1A). Screening was initiated at SMRU until a second screening site was established at IPC, where all unfinished source plates were shipped and assayed. Some plates were assayed more than once in order to obtain a single run with a sufficient Z’ factor of > 0.0 or two moderate-quality runs allowing for identification of reproducibly-active wells (Fig. S1B). Quantification of parasite growth was performed by fixing and staining cultures with recombinant mouse-anti *P. vivax* Upregulated in Infectious Sporozoites 4 (r*Pv*UIS4)^54^. followed by high-content imaging and analysis using an ImageXpress Micro (Molecular Devices) or Lionheart FX (Agilent). Hypnozoites were classified as forms of less than 125 μm^2^ growth area.

#### Normalization, hit selection, and dose-response confirmation in *P. vivax* liver stage assays

Primary screening data were imported into Genedata Screener, Version 15.0.1-Standard and normalized to DMSO (neutral) and inhibitor (monensin) control-treated wells (neutral controls minus inhibitors). For four plates where the monensin control failed due to solubility issues combined with PHH lot variability (Fig. S14), data were normalized using the Robust Z-Score method, which calculates for each well the Robust Z-Score (number of standard deviations off the median) based on the statistics of the compound wells per plate. Genedata multiplicative pattern correction was applied to adjust for plate edge effects. Sixty-two most active (≥ 67% normalized inhibition of hypnozoite numbers) and non-toxic (≤ 40% host cell toxicity) compounds and 10 hydrazinophthalazines were selected for reconfirmation in an 8-point 1:3 dose response following the 8-day protocol with PHH lot BGW using a dose-response of monensin and nigericin as redundant positive controls. Once hydralazine and cadralazine were identified as reconfirmed hits, commercially available batches of powder were obtained (budralazine, Chemcruz cat sc-504334 batch D3019, cadralazine, Chemcruz cat sc-500641 batch B2417, and hydralazine, Selleckchem cat s2562 batch S256202) and used for additional reconfirmation runs using the same 8-day protocol (Fig. 1B-C, S2A).

#### Hit confirmation in *P. cynomolgi* liver stage assays at UGA

*P. cynomolgi* assays at UGA were performed using the step-by-step protocol for the *P. vivax* liver stage assay^36^ with a few modifications. A Japanese macaque (*Macaca fuscata*) was intravenously infected with *P. cynomolgi* Rossan strain cryopreserved ring stage parasites^75^ and allowed to reach patency. When parasitemia reached approximately 5,000 parasites per microliter, *An. dirus* mosquitoes were fed directly on the infected animal over a period of three to four days. The blood-fed mosquitoes were then checked for infection six to eight days by dissecting and staining midguts with 2% mercurochrome to detect oocysts. Two experiments were performed, one with PSH lot CWP, and one with PSH lot NPI. Two days after assay plates (Greiner Bio-One cat 781956) were seeded with 20,000 live PSH per well, sporozoites were dissected from mosquito salivary glands at day 16 post-bloodmeal and allowed to infect cultures. Hits were confirmed using the same 8-day radical cure assay. On treatment days, a pin tool was used to transfer 40 nL of compounds from the source plates to the assay plates. Quantification of *P. cynomolgi* liver stage growth was performed by fixing and staining cultures with 100 ng/mL mouse monoclonal antibody 13.3 (anti-GAPDH) obtained from The European Malaria Reagent Repository (http://www.malariaresearch.eu) followed by high-content imaging and analysis using an ImageXpress Micro (Molecular Devices). Hypnozoites were classified as forms of less than 105 μm^2^ growth area.

#### Hit confirmation in *P. cynomolgi* and *P. vivax* liver stage assays at NITD

Lots of both PSH and PHH were obtained from BioIVT. Hepatocytes were seeded at 22,000 cells per well in a 384-well plate (Corning cat 356667). Prior to and during the infection, the hepatocytes were cultured in BioIVT CP Medium (cat Z990003) with the addition of 1% PSN (Gibco cat 15640055) and 0.1 % gentamicin in the case of *P. vivax*. Two days post-seeding, the hepatocytes were infected with sporozoites dissected from the salivary glands of *An. dirus* mosquitoes. Sporozoites were collected in RPMI 1640 (KD Medical cat CUS-0645). Hepatocytes were infected with 10,000 sporozoites per well and spun for 5 minutes at 200 X *g*. Once the sporozoites were removed after 24 h of incubation, the culture media was exchanged to include 5% PSN in the case of *P. cynomolgi*. On days 4, 5, 6, and 7 post-infection, the hepatocytes received fresh compound addition in media. The cells were fixed on day 8 using 4% paraformaldehyde.

Liver stage parasites were detected by immunofluorescence assay. Hepatocytes were permeabilized for 1 h at room temperature in blocking buffer consisting of 2% Bovine Serum Albumin (Millipore Sigma cat A2153) and 0.2% Triton X-100 (Millipore Sigma cat 648466) in 1X PBS (Gibco cat 20012-027). For *P. cynomolgi* staining, the two in-house primary antibodies used were mouse anti-*Pc*UIS4 monoclonal at 10 ng/mL, and rabbit anti-*Pc*HSP70 polyclonal at 200 ng/mL. For *P. vivax* staining, rabbit anti-*Pv*MIF was used at 1:1000^76^. The primary antibodies were diluted in blocking buffer and incubated overnight at 4 °C. Hepatocytes were washed thrice with 1X PBS and then incubated with secondary antibodies (Invitrogen cat A11013, RRID: AB_2534080 and A11036, RRID: AB_10563566) used at a 1:1000 dilution and Hoechst 33342 (Invitrogen cat H3570) used at 2 μg/mL for 2 h at room temperature. After the incubation, the hepatocytes were washed 3 times with 1X PBS and were stored in 50 μL per well of 1X PBS prior to imaging on an ImageXpress Micro (Molecular Devices).

#### Confirmed hit counterscreens: *P. falciparum* asexual blood stage at Calibr

The SYBR Green I-based parasite proliferation assay^77^ was used to determine the activity of compounds against the asexual blood stage of *P. falciparum* strain Dd2-HLH, a transgenic line expressing firefly luciferase^78^. Briefly, acoustic compound transfer (Labcyte Echo 550) was used to prepare assay ready plates to which parasites in assay medium were added and incubated with compounds for 72 h. SYBR Green I in lysis buffer was used as detection reagent. Fluorescence signal was read on the PHERAstar FSX plate reader (BMG Labtech). Compounds were tested in technical triplicates on different assay plates across three biological replicates performed on different days. Data were uploaded to Genedata Screener, Version 16.0.3-Standard and normalized to DMSO (neutral) and inhibitor control-treated wells (neutral controls minus inhibitors), with 1.25 µM dihydroartemisinin used as a positive control. Dose curves (thirteen point, 1:3 dilution series) were fitted with the four parameter Hill Equation.

#### Confirmed hit counterscreens: *P. falciparum* asexual blood stage at UGA

Budralazine, cadralazine and hydralazine (same catalog and batches as above) were tested using the [^3^H]-hypoxanthine drug susceptibility assay as previously described, with some modifications ^79^. Strain W2^80,81^ was grown in continuous culture using RPMI 1640 media containing 10% heat-inactivated type A+ human plasma, sodium bicarbonate (2.4 g/L), HEPES (5.94 g/L), and 4% washed human type A+ erythrocytes. Cultures were gassed with a 90% N2, 5% O2, and 5% CO2 mixture and incubated at 37 °C. Cultures were sorbitol synchronized to achieve >70% ring stage parasites^82^. Assay were started by establishing a 0.5−0.7% parasitemia and 1.5% hematocrit in complete media. Assays were performed in 96-well plates with a volume of 90 μL/well of parasitized erythrocytes and 10 μL/well of 10X test compound. Dihydroartemisinin was plated as a positive control and DMSO as a negative control. Assay plates were incubated in the above-mentioned gas mixture at 37 °C for 48 h; then, ^3^H-hypoxanthine (185 MBq, Perkin Elmer cat NET177005MC) was added, and plates were incubated for another 24 h. After 72 h of incubation, the assay plates were frozen at −80 °C. Plates were allowed to thaw at room temperature before well contents were collected onto filtermats using a plate harvester (PerkinElmer). A Micro Beta liquid scintillation counter (PerkinElmer) was used to quantify radiation (counts-per-minute) representing relative parasite growth. Values were normalized to controls and plotted using CDD Vault. Potency values represent means of at least two independent experiments.

#### Confirmed hit counterscreens: *P. cynomolgi* asexual blood stage at UGA

Budralazine, cadralazine and hydralazine (same catalog and batches as above) were tested against *P. cynomolgi* DC strain using the [^3^H]-hypoxanthine drug susceptibility assay as previously described, with some modifications^79^. *P. cynomolgi* was grown in continuous culture using RPMI 1640 +GlutaMAX media containing 20% heat-inactivated rhesus serum, hypoxanthine (32 mg/L), HEPES (7.15 g/L), additional glucose (2g/L), and 5% washed rhesus erythrocytes. Cultures were incubated at 37°C under mixed gas conditions of 90% N2, 5% O2, and 5% CO2. Schizonts were synchronized over a 60/20 Percoll gradient to achieve >90% late-stage parasites. Assays were started the following day when ring-stage parasites were present. Parasites were prepped for assay by establishing 0.5% ring-stage parasitemia and 2% hematocrit in complete media without hypoxanthine. Assays were performed in 96-well plates with a volume of 90 μL/well of parasitized erythrocytes and 10 μL/well of 10X test compounds. Compounds were plated from a starting concentration of 5 μM in an 11-point 1:2 dilution series and tested in duplicate. Uninfected RBCs were plated as a positive control and DMSO was used as a negative control. ^3^H-hypoxanthine (185 MBq, Perkin Elmer cat NET177005MC) was then added to all wells and plates were incubated under the previously-mentioned conditions for 72 h. After 72 h the assay plates were frozen at −80 °C. Plates were thawed the following day at room temperature and well contents were collected onto filtermats using a plate harvester (PerkinElmer). A Micro Beta liquid scintillation counter (PerkinElmer) was used to quantify radiation (counts-per-minute) representing relative parasite growth. Values were normalized to controls and plotted using CDD Vault. Potency values represent means of at least two independent experiments.

#### Confirmed hit counterscreens: *P. berghei* liver stage at Calibr

For *P. berghei* liver stage assays, a colony of *An. stephensi* mosquitoes was maintained in the UGA Sporocore using methods previously described^83^. In summary, adults were fed 5% dextrose (w/v) and 0.05% para-aminobenzoic acid (w/v) soaked into cotton pads and kept at a temperature of 27 °C, relative humidity of 75-85%, and a 12 h light/dark cycle. PbGFP-LUC_CON_ sporozoites were produced as previously described^83^. In summary, female C57BL/6 or Hsd:ICR(CD-1) mice (Envigo) were injected intraperitoneally with 5×10^6^ - 5×10^7^ blood stage parasites in 500 μL PBS 3-4 days before mosquito infections. Once parasitemia reached 2-6%, mice were anesthetized with 0.5 mL 1.25% 2,2,2-Tribromoethanol (v/v, Avertin, Sigma-Aldrich) and placed on top of cage of *An. stephensi* mosquitoes (3-7 days post-emergence) for 20 min to serve as an infectious bloodmeal. Infected mosquitoes were shipped to Calibr, where sporozoites were dissected out of mosquito salivary glands and used for luciferase-based infection assay as previously described^84^. Briefly, HepG2 cells (ATCC cat HB-8065, RRID: CVCL_0027) were infected with freshly dissected sporozoites. The infected cells were incubated with compounds of interest in 1536-well plates for 48 hours, and intracellular parasite growth was measured using bioluminescence. Compounds were tested in technical triplicates on different assay plates across three biological replicates performed on different days. Data were uploaded to Genedata Screener, Version 16.0.3-Standard and normalized to DMSO (neutral) and inhibitor control-treated wells (neutral controls minus inhibitors), with 1 µM KAF156 used as a positive control. Dose curves (thirteen point, 1:3 dilution series) were fitted with the four parameter Hill Equation.

#### Confirmed hit counterscreens: Mammalian cell cytotoxicity at Calibr

HepG2 (ATCC cat HB-8065, RRID: CVCL_0027) and HEK293T (ATCC cat CRL-3216, RRID: CVCL_0063) mammalian cell lines were maintained in Dulbecco’s Modified Eagle Medium (DMEM, Gibco) with 10% heat-inactivated HyClone FBS (GE Healthcare Life Sciences), 100 IU penicillin, and 100 µg/mL streptomycin (Gibco) at 37 °C with 5% CO_2_ in a humidified tissue culture incubator. To assay mammalian toxicity of hit compounds, 750 HepG2 and 375 HEK293T cells/well were seeded, respectively, in assay media (DMEM, 2 % FBS, 100 U/mL penicillin, and 100 µg/mL streptomycin) in 1536-well, white, tissue culture-treated, solid bottom plates (Corning cat 9006BC) that contained acoustically transferred compounds in a three-fold serial dilution starting at 40 µM. After a 72 h incubation, 2 µL of 50% Cell-Titer Glo (Promega cat G7573) diluted in water was added to the cells and luminescence measured on an EnVision Plate Reader (PerkinElmer).

#### Additional ReFRAME hit confirmation using an improved *P. vivax* liver stage assay

Twelve hits were re-confirmed using the 12-day radical cure assay, implementing three assay improvements^36^. First, 100 μM 1-ABT (Caymen Chem cat 15252) was added to media on treatment days to reduce hepatic metabolism. Second, the assay endpoint was extended 4 days to allow for nonviable liver stage forms to be cleared from cultures and therefore not be quantified during high-content imaging. Third, nigericin replaced monensin as the positive ionophore control. Confirmation was performed with one independent experiment for all compounds except cadralazine, which was confirmed in four independent experiments.

#### Epigenetic inhibitor library screen against *P. vivax* liver stages

The Epigenetic Inhibitor library (Targetmol, cat L1200), containing 773 compounds at 10 mM, was purchased and re-plated in pintool-ready 384-well source plates with 200 μM nigericin and DMSO control wells. The library was screened using the 12-day radical cure assay noted above. The 24 hits exhibiting the highest inhibition against hypnozoites were replated in a dose-response for confirmation of activity in a 12-day radical cure assay as described above. Confirmation was performed with two independent experiments. The ACS inhibitors MMV019721 and MMV084978 were kindly provided by MMV and tested in dose-response in a 12-day radical cure assay as described above. Potency was determined from four independent experiments.

#### Assessment of effect of 1-ABT on hepatic cytochrome P450 3A4 activity

Two experiments were performed, one on uninduced PHHs, and another on rifampicin-induced PHHs (BioIVT, lot BGW). Cells were thawed and 18,000 live cells/well were seeded into collagen-coated 384-well plates as described above. Media was exchanged every-other-day until day 7 post-seed when media exchange included a dilution series of 1-ABT. One hour after addition of 1-ABT, cytochrome P450 3A4 activity (CYP3A4) was measured using a luciferin-IPA kit (Promega cat V9001) following the lytic protocol with 3 μM IPA. Lysed well contents were transferred to a white 384-well luminometer plate (Greiner Bio-One cat 201106) before reading on a Spectramax i3X (Molecular Devices) with a 1 sec integration time. In the second experiment, cells were similarly seeded and cultured before addition of 25 μM rifampicin (MP Biomedial cat BP2679-250), or an equivalent v/v DMSO vehicle control, in media on days 4 and 6. At day 7 post-seed, CYP3A4 activity was measured following addition of 1-ABT as above. The fold change was calculated between induced and uninduced wells at each 1-ABT dilution point.

#### Assessment of effect of 1-ABT on hepatic metabolism using mass spectrometry

Primary human hepatocytes (lot BGW, BioIVT) were thawed and 18,000 live cells/well were seeded into collagen-coated 384-well plates as described above. Media was exchanged every-other-day until day 7 post-seed when cells were treated with 100 μM 1-ABT, or an equivalent v/v vehicle control, in media for 1 h. Cells were then incubated with standard substrates for characterization of phase I and II hepatic metabolism, including: 30 μM 7-hydroxycoumarin (UGT/ST), 40 μM coumarin (CYP2A6), 500 μM chlorzoxazone (CYP2E1), 50 μM dextromethorphan (CYP2D6), 24 μM midazolam (CYP3A4/5), 500 μM S-mephenytoin (CYP2C19), 600 μM testosterone (CYP3A4), 1 mM tolbutamide (CYP2C9), 500 μM phenacetin (CYP1A2), or 400 μM bupropion (CYP2B6). The reaction was stopped at 1 h by addition of an equal volume of ice cold methanol. Metabolite formation was quantified using UPLC-MS/MS or LC-MS/MS (7-HC, 7-HCS, and 7-HCG). Samples were thawed, vortexed, and centrifuged for 5 min at 5000 rpm. Standards, controls, blanks, and study samples were added to an HPLC autosampler vial and injected into the UPLC-MS/MS or LC-MS/MS systems. Analyses were run using an Acquity UPLC (Waters) or Agilent 1100 HPLC (Agilent) and Quattro premier XE (Waters) or Quattro Premier ZSpray (Waters) mass spectrometers. Quantification was performed using a quadratic least squares regression algorithm with 1/X^2^ weighting, based on the peak area ratio of substrate or metabolite to its internal standard. Metabolite formation rate was calculated as pmol/min/10^6^ cells.

#### Combination drug studies in *P. vivax* liver stages

Powders of cadralazine (same batch as above), 5-azacytidine (Caymen Chem, cat 11164), and nigericin, were diluted to 50 mM, 50mM, and 200 μM, respectively, in DMSO, before being diluted to 100 μM, 100 μM, and 400 nM, respectively, in hepatocyte culture media (BioIVT, cat Z99029). Cadralazine and 5-azacytidine were then plated in the first column of two 96-well plates at volumetric ratios of 1:0, 8:1, 6:1, 4:1, 2:1, 1:1, 1:2, 1:4, 1:6, 1:8, and 0:1 such that the net volume per well was 200 μL (nigericin and DMSO controls were also diluted as such). Each mixture was then diluted in a 12-point, 2-fold dilution series by mixing 100 μL of mixture to 100 μL media in subsequent columns using a multichannel pipettor. A 384-well *P. vivax* liver stage assay plate was started using the 12-day protocol as above, and on day 5, 6, and 7 post-infection, media was removed from the 384-well plate using the inverted spin method^36^ followed by addition of 20 μL of fresh media. Then, a multichannel pipettor was used to transfer 20 μL of the mixtures (made fresh daily) from the 96-well dilution series plates to the 384-well plates, thereby establishing a highest 1:0 and 0:1 treatment dose of 50 μM. The assay was fixed, stained, imaged, and parasite growth quantified as described above. Parasite growth data were normalized to the DMSO control and loaded into Prism (Graphpad) for curve fitting using the setting “log(inhibitor) vs. response -- variable slope (four parameters) least squares fit.” The EC_50_s of each ratio were used to calculate Fractional Inhibitory Concentrations (FICs) and plot isobolograms as previously described^85^.

#### Immunofluorescent staining of methyl-cytosine modifications in *P. vivax* liver stages

Sporozoites from three different *P. vivax* cases were infected into PHH lot BGW at day 2 post-seed (for case 1) or day 3 post-seed (for cases 2 and 3) in 384-well plates (Greiner Bio-One cat 781956) using the same methods for initiating *P. vivax* liver stage screening assays described above. Cultures were fixed at day 6 post-infection and stained with r*Pv*UIS4 and Hoechst 33342 as previously described^36,54^. Cultures were then stained for either with rabbit anti-5mC monoclonal antibody (clone RM231, ThermoFisher Scientific cat MA5-24694, RRID: AB_2665309) or rabbit anti-5hmC monoclonal antibody (clone RM236, ThermoFisher Scientific cat MA5-24695, RRID: AB_2665308) using methods adapted from those previously described^51^. In summary, cultures were re-permeablilized with 0.1% (v/v) Triton-X 100 for 20 min at room temperature and then washed thrice with 1X PBS. Chromatin was then denatured with 4 N HCl for 30 min at room temperature and washed thrice with 1X PBS. The denaturing reaction was then neutralized with 100 mM Tris (pH 8.0) for 10 min at room temperature and washed thrice with 1X PBS. Cultures were then quenched with 50 mM NH_4_Cl for 10 min at room temperature and washed thrice with 1X PBS. Cultures were then blocked with 0.1% (v/v) Tween 20 and 2% (w/v) Bovine Serum Albumin for 10 min at room temperature and washed thrice with PBS. Cultures were then stained with either antibody diluted to 10 μg/mL in PBS overnight at 4 °C and washed thrice with 1X PBS. Cultures were then stained with 10 μg/mL Texas Red-conjugated, goat anti-rabbit IgG secondary antibody (ThermoFisher Scientific, cat T-2767, RRID: AB_2556776) overnight at 4 °C and washed thrice with 1X PBS. For a negative stain control, a separate set of infected wells were prepared as above and stained with secondary antibody only (2’ control, Fig. 3B). High resolution images of individual parasites and PHH nuclei were obtained by capturing 8 planes in the Z dimension using a 100X objective on Deltavision Core (GE Healthcare Life Sciences) and deconvoluted using softWoRx (GE Healthcare Life Sciences) (Fig. 3A, S6-8). An ImageXpress Micro high content imager was used to quantify methyl-cytosine modifications for the entire population of parasites from each case. A 20X objective was used to capture 25 fields of view from each well (covering the entire growth area) of the 384-well plate. Using the associated MetaXpress high content analysis software, the r*Pv*UIS4 stain from each parasite was used to define parasite objects, and the 5mC or 5hmC staining of host cell nuclei were used to define positive methyl-cytosine modification objects. The 2-dimensional area of intersection of both objects was then quantified for each parasite and forms less than 125 μm^2^ were quantified as hypnozoites (Fig. 3B).

#### Immunofluorescent staining of methyl-cytosine modifications in *P. cynomolgi* liver stages

Japanese macaques (*Macaca fuscata*) were intravenously infected with *P. cynomolgi* M/B strain^86^ and allowed to reach patency before skin feeding to *An. dirus* mosquitoes as described above. One round of macaque infection, mosquito dissection, and culture infection was performed with PSH lot NPI and a second round was performed with PSH lot NNF. Two days after assay plates (Greiner Bio-One cat 781956) were seeded with 20,000 primary simian hepatocytes per well, sporozoites were dissected from mosquito salivary glands at day 16 post-bloodmeal and allowed to infect cultures. Cultures were fixed on day 8 (experiment 1) or 12 (experiment 2) post-infection and stained for 5mC and 5hmC as described above. An ImageXpress Micro high content imager was used to quantify methyl-cytosine modifications for the entire population of *P. cynomolgi* liver stage parasites. A 20X objective was used to capture 25 fields of view from each well (covering the entire growth area) of the 384-well plate. Using the associated MetaXpress high content analysis software, the GAPDH stain from each liver stage parasite was used to define parasite objects, and the 5mC or 5hmC staining of host cell nuclei were used to define positive methyl-cytosine modification objects. The 2-dimensional area of intersection of both objects was then quantified for each parasite and forms less than 105 μm^2^ were categorized as hypnozoites (Fig. S10).

#### Collection of *P. vivax* and *P. cynomolgi* sporozoites for methyl-cytosine characterization

For quantification of 5mC modification levels by mass spectrometry, sporozoites from 3 different *P. vivax* cases, numbering 18.7×10^6^ from case 1, 10×10^6^ from case 2, and 14.7×10^6^ from case 3, were dissected from infected *An. dirus* mosquitoes at IPC as previously described^36^ and cryopreserved as previously described^87^. For quantification of DMNT activity from nuclear extracts, sporozoites from 2 different *P. vivax* cases, numbering 21×10^6^ from case 1 and 20×10^6^ from case 2, were similarly dissected and cryopreserved. To serve as a negative control, salivary glands from uninfected mosquitoes at IPC were similarly dissected and cryopreserved. A total of 4.8×10^6^ sporozoites for mass spec and 34.1×10^6^ sporozoites for DNMT activity assays were also collected from *An. dirus* mosquitoes infected from feeding on a rhesus macaque infected with *P. cynomolgi* M/B strain at ENPRC and cryopreserved as described above. To serve as a negative control, salivary glands and ovaries from uninfected mosquitoes at ENPRC were similarly dissected and cryopreserved. For mapping of methyl-cytosine modifications by bisulfite sequencing, sporozoites from three different *P. vivax* cases, numbering 9.8×10^6^ from case 1, 12.3×10^6^ from case 2, and 15.1×10^6^ from case 3, were dissected from infected *An. dirus* mosquitoes at IPC and cryopreserved as described above. A total of 5.3×10^6^ sporozoites were also collected from *An. dirus* mosquitoes infected from feeding on a rhesus macaque infected with *P. cynomolgi* M/B strain at ENPRC and cryopreserved as described above. Frozen sporozoites and salivary glands were shipped from IPC and ENPRC to University of California, Riverside on dry ice.

#### Quantification of 5mC, 5hmC and 2’-deoxyguanosine (dG) in genomic DNA by LC-MS/MS/MS

Parasite pellets were lysed with 100 µl lysis buffer (20 mM Tris, pH 8.1, 20 mM EDTA, 400 mM NaCl, 1% SDS and 20 mg/ml proteinase K) and incubated at 55 ^0^C overnight. Saturated solution of NaCl (0.5X volume of reaction mixture) was subsequently added to the digestion mixture and incubated at 55 °C for another 15 min. The samples were centrifuged at 14500 RCF for 30 min at 4 °C and the supernatant removed to a 1.5-mL microcentrifuge, genomic DNA (gDNA) was then precipitated with 2X volume of 100% chilled ethanol and resuspended in 95 μL water. Samples were then treated with 3 μL of 10 mg/mL RNase A and 2 μL of 25 units/μL RNase T1 and incubated overnight at 37 °C. gDNA was then extracted by chloroform/isoamyl alcohol solution, precipitated again with 100% chilled ethanol, and washed with 70% ethanol. The gDNA pellets were then dissolved in nuclease-free water. One μg of gDNA was enzymatically digested into mononucleosides using nuclease P1 and alkaline phosphatase. Enzymes in the digestion mixture were removed by chloroform extraction. The resulting aqueous layer was dried by using a SpeedVac and the dried residues were subsequently reconstituted in doubly distilled water. Approximately 5 ng of the DNA digestion mixture was injected for LC-MS/MS/MS analyses for quantifications of 5mC, 5hmC and dG. An LTQ XL linear ion-trap mass spectrometer equipped with a nano electrospray ionization source and coupled with an EASY-nLC II system (Thermo Fisher Scientific) was used for the LC-MS/MS/MS experiments. The amounts of 5mC, 5hmC and dG (in moles) in the nucleoside mixtures were calculated from area ratios of peaks found in the selected-ion chromatograms for the analytes over their corresponding isotope-labeled standards, the amounts of the labeled standards added (in moles), and the calibration curves. The final levels of 5mC and 5hmC, in terms of percentages of dG, were calculated by comparing the moles of 5mC and 5hmC relative to those of dG.

#### Extraction of nuclear protein extracts

Cryopreserved sporozoites, or parasites extracted from red blood cells by saponin lysis, were resuspended in 1 ml of cytoplasmic lysis buffer (20mM HEPES pH 7.9, 10 mM KCl, 1mM EDTA, 1mM EGTA, 1mM dithiothreitol (DTT), 0.5 mM AEBSF, 0.65% Igepal, 1X Roche complete protease inhibitor cocktail) and incubated for 10 min on ice. Nuclei were separated from cytoplasmic fraction by 10 min of centrifugation at 1500 RCF followed by two washes with cytoplasmic lysis buffer and one time wash with ice cold 1x PBS. Nuclei pellets were resuspended in 100 µl of nuclei lysis buffer (20 mM HEPES pH 7.9, 0.1 M NaCl, 1mM EDTA, 1 mM EGTA, 1mM DTT, 25 % glycerol, 0.5 mM AEBSF, 1X Roche complete protease inhibitor cocktail) for 20 min at 4 °C with rotation. Nuclear extracts were cleared by 10 min of centrifugation at 6000 RCF. Protein concentration of nuclear extract was quantified by BCA assay and DNMT assays were performed immediately after estimation of protein concentration.

#### DNA methyltransferase assay

DNMT activity of nuclear extracts from *P. cynomolgi* sporozoites, *P. vivax* sporozoites, and uninfected mosquito salivary glands were measured using the Epiquik DNMT activity/inhibition assay ultra-kit (cat P-3010) following the manufacturer’s instructions. Purified bacterial DNMT enzyme was used as a positive control. A blank control was used to subtract the residual background values. Each reaction was performed in duplicate. DNMT activity was measured in relative unit fluorescence per h per mg of protein for 10 min at 1 min intervals.

#### Bisulfite conversion and library preparation

*P. cynomolgi* and *P. vivax* sporozoites were lysed using 100 µl of lysis buffer containing 20mM Tris (pH 8.1), 20 mM EDTA, 400 mM NaCl, 1 % SDS (w/v) for 30 min at room temperature followed by addition of 20 µl of proteinase K (20 mg/ml) to the pellet and incubated at 55 °C overnight. The gDNA mixture was purified with phenol-chloroform followed by chloroform. Precipitation of gDNA was performed using chilled ethanol and treated with RNase A followed by another round of ethanol precipitation. 50 ng of unmethylated lambda DNA was added as a control to each sample before bisulfite conversion of the DNA. 500 ng of gDNA of each sample was used for the bisulfite conversion following the manufacturer’s instructions (Epitect fast bisulfite conversion kit, Qiagen cat 59824). Libraries from bisulfite converted DNA were prepared using the Accel-NGS methyl-Seq DNA library kit (Swift biosciences cat 30024). Libraries were generated following the manufacturer’s instructions and DNA was cleaned through SPRI select beads (Beckman Coulter). Libraries were sequenced using the NOVASeq platform.

#### DNA methylation analysis

Four sets of reads for *P. vivax* and *P. cynomolgi* were analyzed. Read qualities were checked with FastQC v0.11.8. FastQC indicated the presence of adapter contamination and overrepresented k-mers. As a result, (1) the first 9-14 base pairs were trimmed and (2) reads with overrepresented k-mers were discarded (see Fig. S11C for summary statistics after the cleaning step). Reads were mapped against the corresponding reference genomes downloaded from PlasmoDB (namely, PlasmoDB-48_Pfalciparum3D7, PlasmoDB-48_PcynomolgiB and PlasmoDB-48_PvivaxP01) using Bismark v0.22.2 with default parameters. To determine the bisulfite conversion rate, reads were also mapped against the lambda phage (see Fig. S11C for the conversion rate). Alignment files for the replicates were merged together using Samtools v1.9. Read methylation levels were obtained using Bismark v0.22.2 with default parameters (see Fig. S11C).

A cytosine in the genome was considered methylated if (1) the number of reads covering that cytosine was higher than a given threshold (10 for *P. falciparum*, 5 for *P. vivax* and 3 for *P. cynomolgi*) and (2) the ratio of methylated reads over all reads covering a cytosine was higher than a given threshold (we chose 0.1 for this second threshold). Genome-wide cytosine density and methylated cytosine density in Fig. 4A-B were calculated in 1 kbp non-overlapping sliding windows using a custom script (available at https://github.com/salehsereshki/pyMalaria). The distribution of CG, CHG, CHH methylation in Fig. 4C-D were obtained by computing the number of methylated cytosines in each context over all the methylated cytosines. For the methylation analyses in genes in Fig. 4E, (1) 500 bp flanking regions and gene body were split into five bins, (2) methylation levels were averaged across all the genes using a custom script (available at the https://github.com/salehsereshki/pyMalaria). To study the correlation between cytosine methylation and gene expression, the same gene body computation was done for the 10% high and low expressed genes using a previously reported *P. vivax* transcriptome^32^. These plots are represented in Fig. 4F.

### Supplementary Text

#### Immunofluorescent staining of 5mC and 5hmC in *P. vivax* blood stage parasites

An immunofluorescent staining approach has been used to detect both 5mC and 5hmC in *P. falciparum* blood stage parasites^51^, thus we sought to confirm these marks in *P. vivax* blood stages. *P. vivax* blood samples were collected between 2017 and 2019 by active and passive case detection from individuals residing in Mondulkiri, Eastern Cambodia. The presence of *P. vivax* was determined using an RDT (CareStartTM Malaria Pf/pan RDTs, Accessbio) or microscopy and monoinfections were confirmed by RT-PCR using species-specific primers^88^. Venous blood used was collected in lithium heparin tubes and immediately processed on-site in a mobile laboratory. Leukocytes were depleted using NWF filters^89^. The leukocyte depleted parasitized red blood cells were cryopreserved using glycerolyte 57 solution (Baxter) and immediately stored in liquid nitrogen^90^. Blood isolates were thawed by addition of 12%, then 1.6%, and then 0.9% (w/v) NaCl solution followed by heparin treatment for 10 min at 37 °C. Blood stage parasites were then purified from thawed isolates using a KCl-Percoll density gradient^91^ followed by a wash with RPMI and two washes with 1X PBS. Parasites were then fixed with 3% (v/v) paraformaldehyde and 0.01% (v/v) glutaraldehyde in 1X PBS for 1 h at 4 °C. After fixation, blood stage parasites were permeabilized, denatured, neutralized, quenched, and blocked as described above. Staining for 5mC and 5hmC was carried out as described above except the primary and secondary antibodies were diluted to 1 μg/mL instead of 10 μg/mL. Parasites were stained with 10 μg/mL Hoechst 33342 for 30 min at room temperature and then washed twice with 1X PBS after staining. Parasites were mounted on a coverslip and imaged with a 100X objective on a Leica DM250. While we did detect 5mC and 5hmC methylation in residual human white blood cells, we could not confirm positive 5mC or 5hmC staining in *P. vivax* blood stage parasites from these isolates (Fig. S18). These negative results could be due to one or more factors. First, while *P. falciparum* blood stage cultures can reach parasitemias above 10%, *P. vivax* blood stages cannot be propagated *in vitro* and the parasitemia of isolates is typically just above the level of detection. Second, *P. vivax* blood stage isolates were cryopreserved before staining and the stability of DNA methylation after cryopreservation is unknown. Third, the hydrochloric acid treatment needed to denature chromatin during the stain protocol causes red cells to aggregate, thereby making finding and imaging *P. vivax* blood stages difficult.

**Fig. S1.**
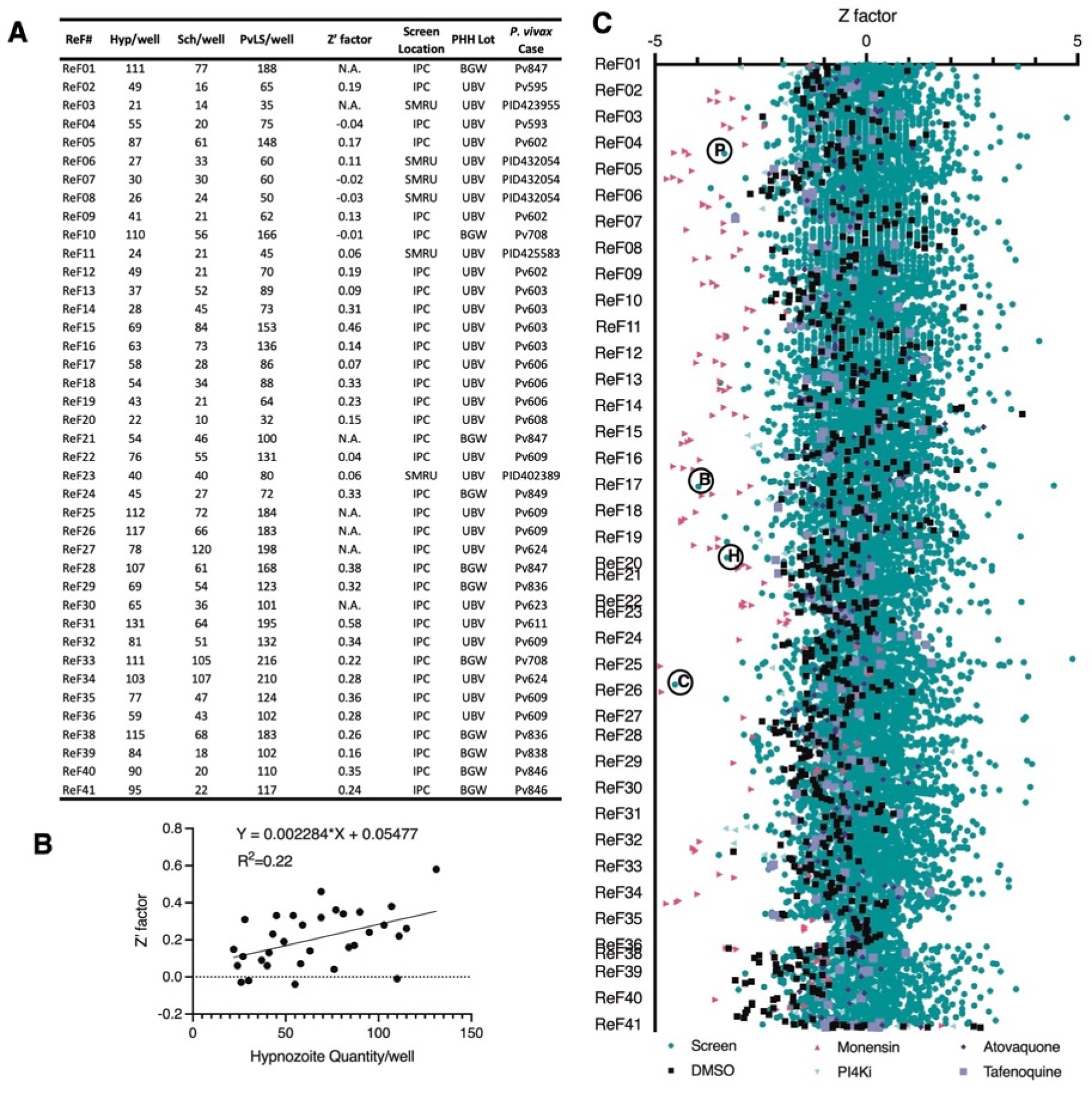
ReFRAME screen run details. (**A**) Table of each ReFRAME plate (40 plates labelled 1-41, with 37 skipped) run metrics including average hypnozoites and schizont counts per well, Z’ factor for 1 μM monensin wells, screening location (Shoklo Malaria Research Unit, Thailand, or Pasteur Institute of Cambodia) PHH lot used, and *P. vivax* patient isolate used. Due to an error during library plating, some plates contained only 1 well of monensin, preventing calculation of a Z’ factor for those plates (listed as N.A.). (**B**) Simple linear regression correlating Z’ factor with average hypnozoite count per well. (**C**) Index chart from Fig. 1A with phosphatidylinositol 4-kinase inhibitor (PI4Ki) KDU691 or MMV390048, tafenoquine, and atovaquone controls added. Teal circle: library, black square: DMSO, pink triangle: 1 μM monensin, light green inverted triangle: 1 μM P4Ki, black diamond: 1 μM atovaquone, purple square: 10 μM tafenoquine. Some hits discussed in this report are noted with black circles; P: poziotinib, B: budralazine, H: hydralazine, C: cadralazine.

**Fig. S2.**
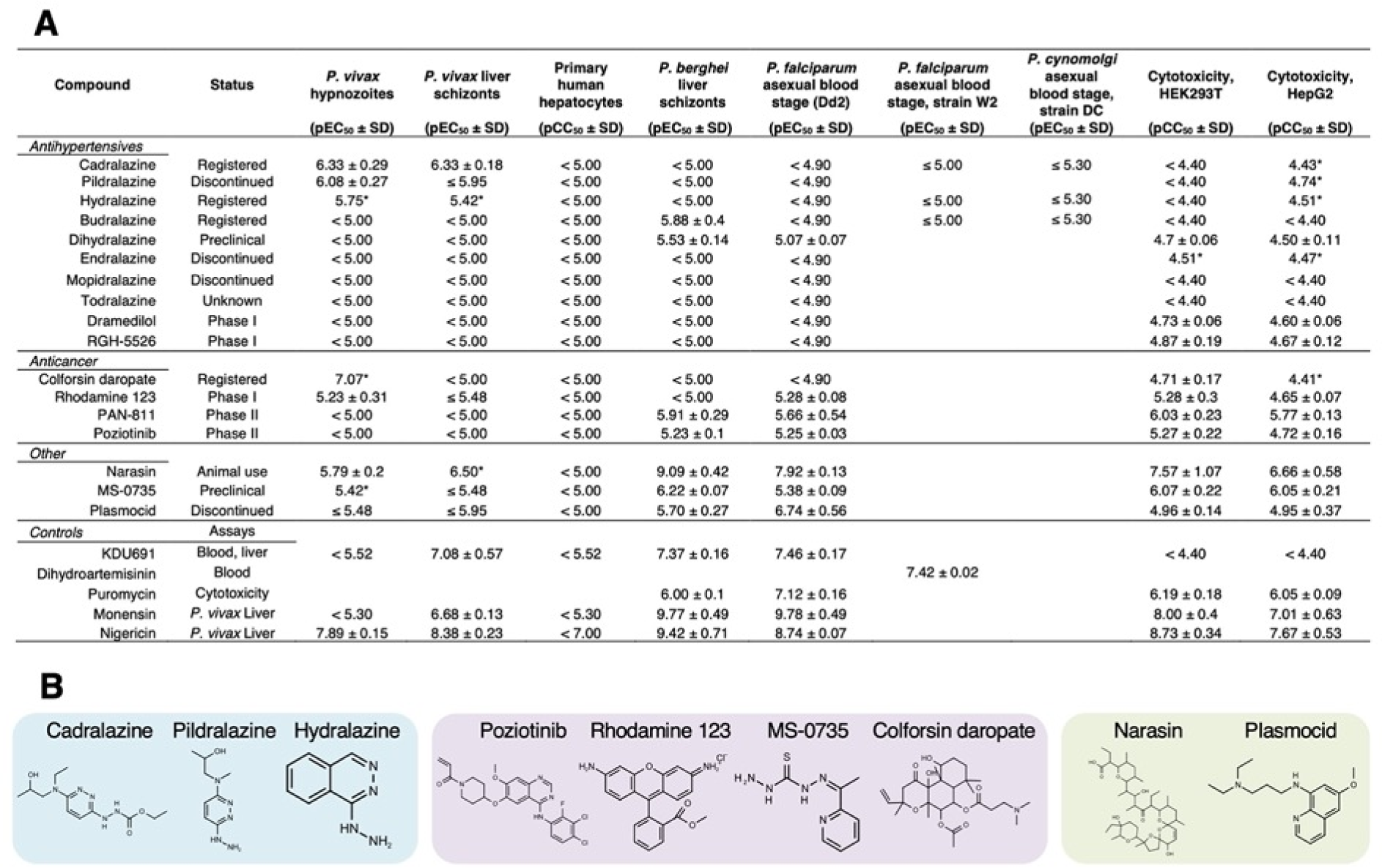
Dose-response confirmation and counterscreens of primary screen hits and analogs. (**A**) Primary screen hits and structurally- or mechanistically-related compounds were tested by dose-response in 8-day *P. vivax* liver stage assays at Institute Pasteur of Cambodia and counterscreened against *P. berghei* liver schizonts, *P. falciparum* asexual blood stages, *P. cynomolgi* asexual blood stages, HEK293T, and HepG2. Values represent pEC_50_ or pCC_50_ ± SD of all independent experiments (n=2-6) for which a pEC_50_ or pCC_50_ was obtained. An asterisk (*) indicates only one independent experiment resulted in a calculated pEC_50_ or pCC_50_. (**B**) Structures of hits which confirmed to be active against *P. vivax* hypnozoites in dose-response assays; blue: hydralazine analogs, purple: other novel hits, green: re-discovery of compounds previously demonstrated to have hypnozonticidal activity *in vitro* or antirelapse activity *in vivo*.

**Fig. S3.**
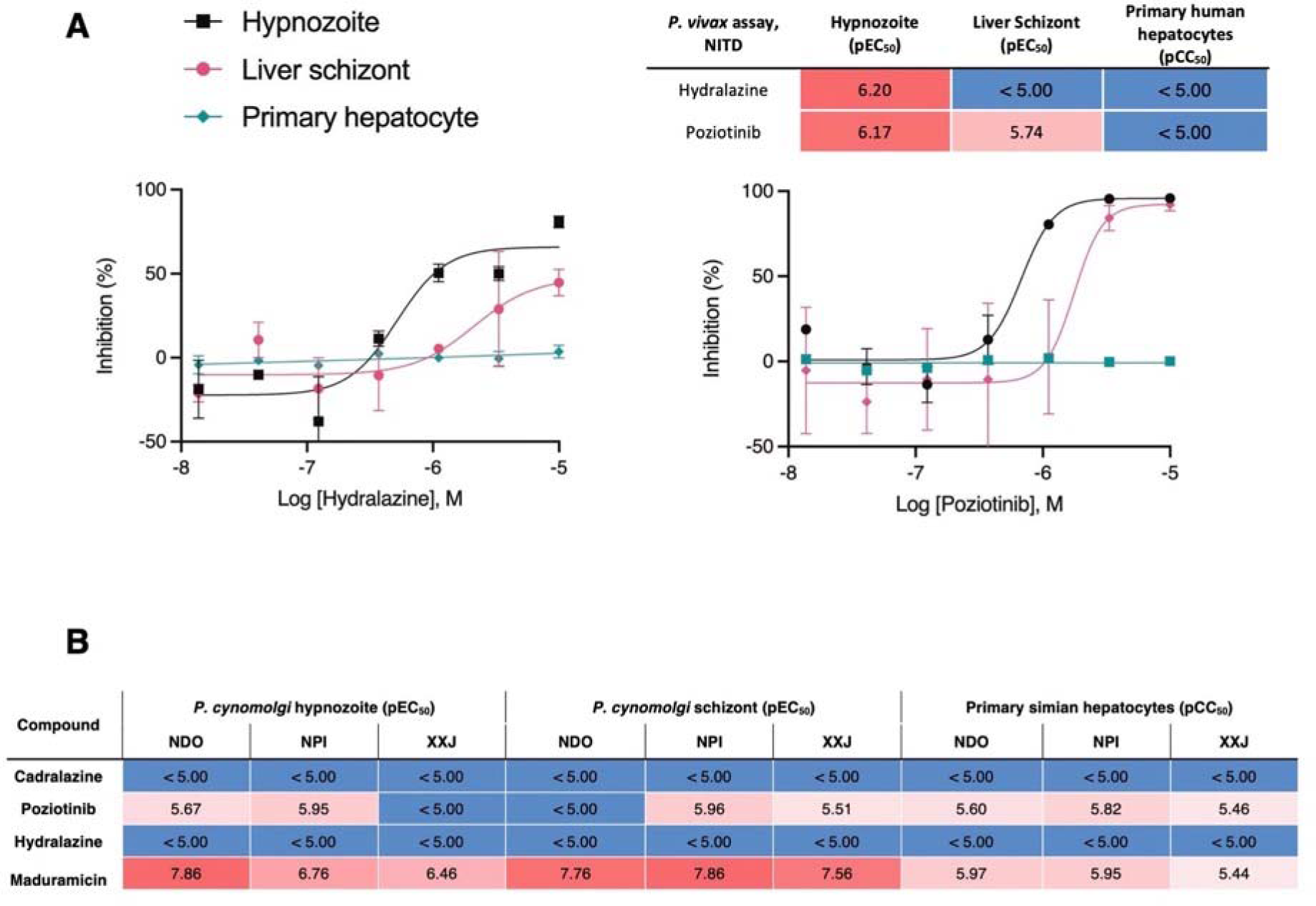
Select ReFRAME hits confirmed at Novartis Institute for Tropical Diseases (NITD). (**A**) Dose-response curves for hydralazine and poziotinib against *P. vivax* liver forms assayed at NITD. All replicate wells were plotted together from a single independent experiment, bars represent SEM. (**B**) Potency data (pEC_50_) for ReFRAME hits against *P. cynomolgi* liver forms assayed at NITD in primary simian hepatocyte (PSH) lots NDO, NPI, XXJ infected with one batch of *P. cynomolgi* sporozoites. Cytotoxicity (pCC_50_) against PSH was measured using nuclei counts. Maduramicin is a positive control with activity against *P. cynomolgi* hypnozoites (A,B) Heat maps represent red as more potent and blue as inactive at highest dose tested.

**Fig. S4.**
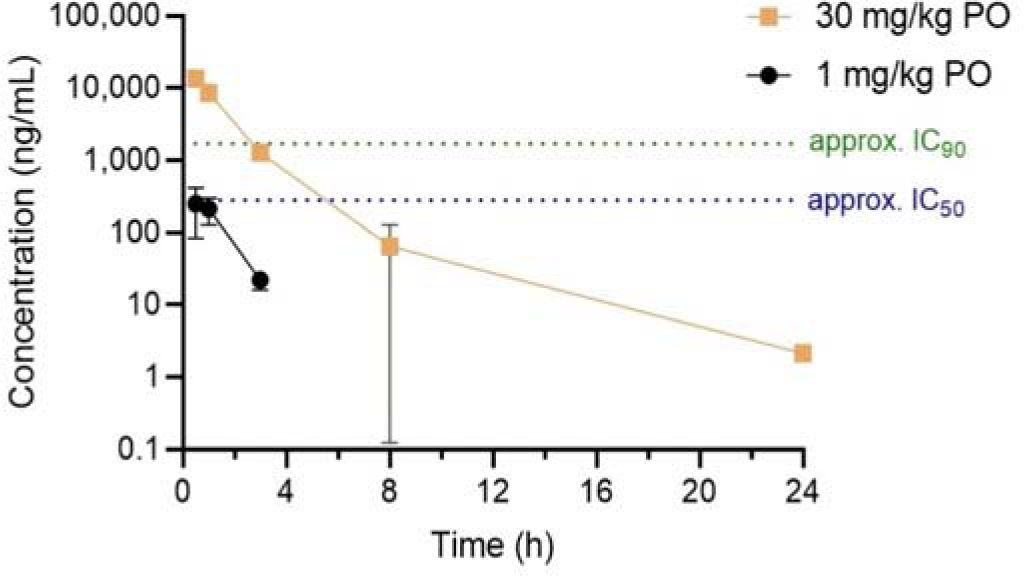
Pharmacokinetics of cadralazine in nonhuman primates. Mean plasma concentration of cadralazine was measured in three males rhesus macaques after oral dosing. Plasma was collected following a 1 mg/kg dose, and again following a 30 mg/kg dose. Bars represent SD. The approximate IC_50_ and IC_90_ from *P. vivax* hypnozoite assays are indicated.

**Fig. S5.**
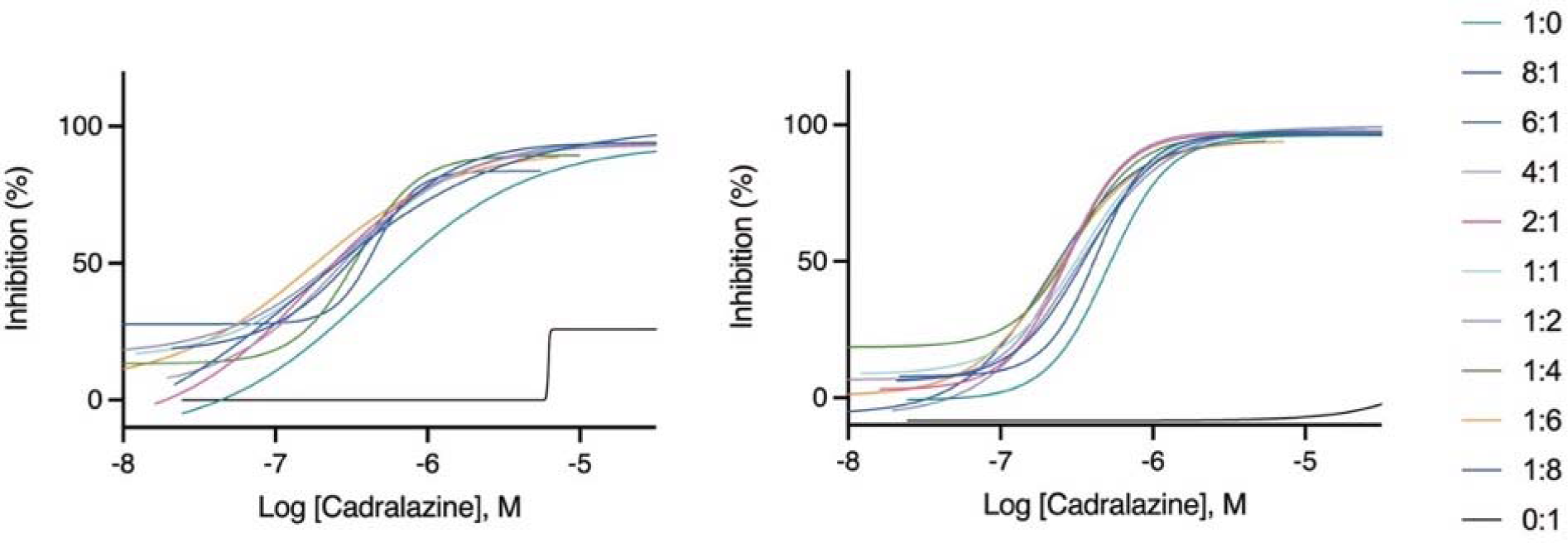
Synergistic effect of cadralazine and 5-azacytidine in *P. vivax* liver stage assays. Dose-response curves for cadralazine with all fixed ratios of 5-azacytidine against *P. vivax* hypnozoites. Cadralazine alone is represented as 1:0, 5-azacytidine alone is represented as 0:1 and plotted on the cadralazine chart for comparison. Left and right charts represent two independent experiments.

**Fig. S6.**
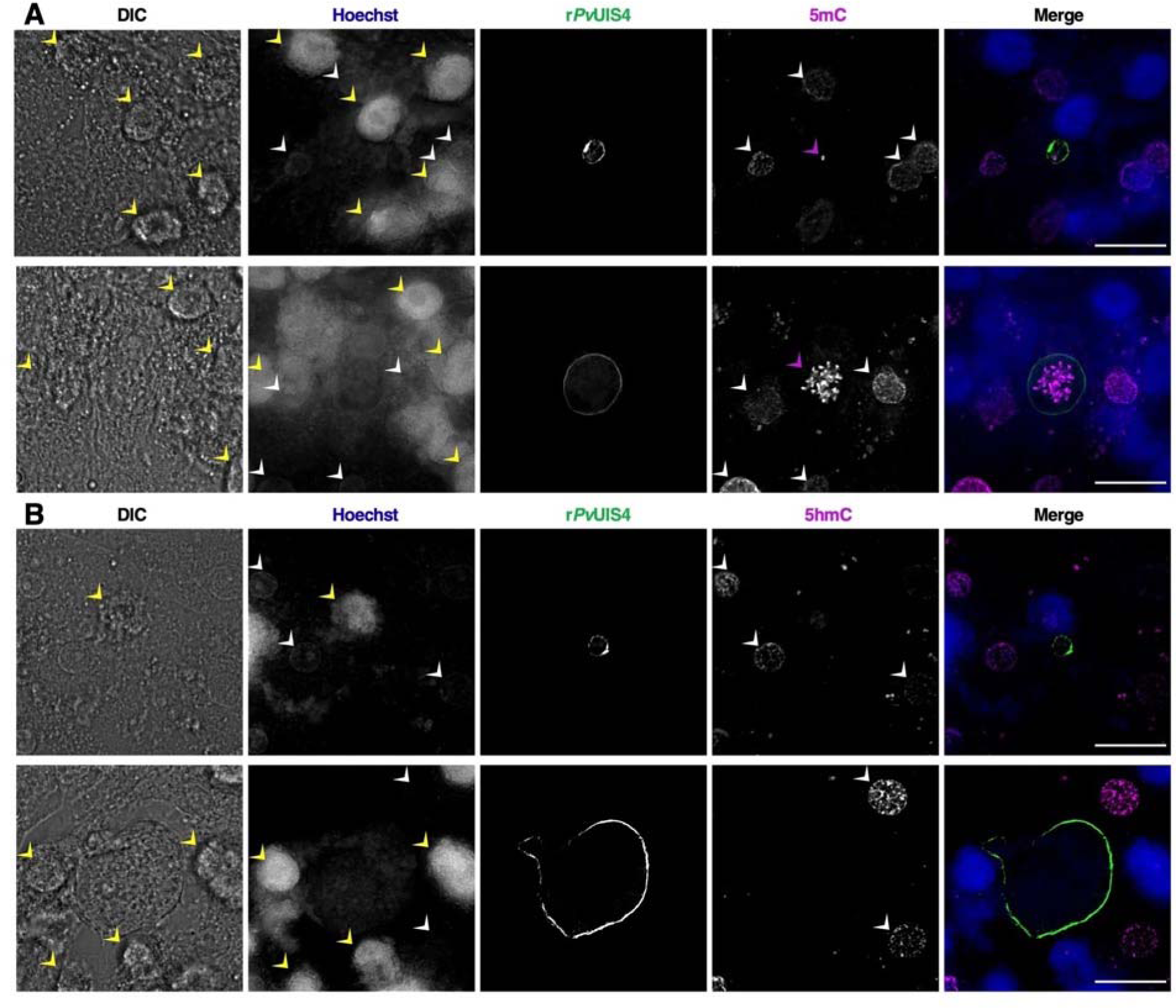
Cytosine modifications in *P. vivax* liver forms, full panels from Case 1 (also shown in Fig. 3). (**A**) Immunofluorescent imaging of a 5mC-positive *P. vivax* hypnozoite (top) and schizont (bottom) at day 6 post-infection. (**B**) Immunofluorescent imaging of a 5hmC-negative *P. vivax* hypnozoite (top) and schizont (bottom) at day 7 post-infection. Yellow arrows indicate autofluorescence in the blue channel associated with cell debris above the hepatocyte monolayer. White arrows indicate hepatocyte nuclei which are dimly-stained with Hoechst 33342 and positive for 5mC or 5hmC. Purple arrows indicate 5mC-positive foci within the parasite. Bars represent 20 µm.

**Fig. S7.**
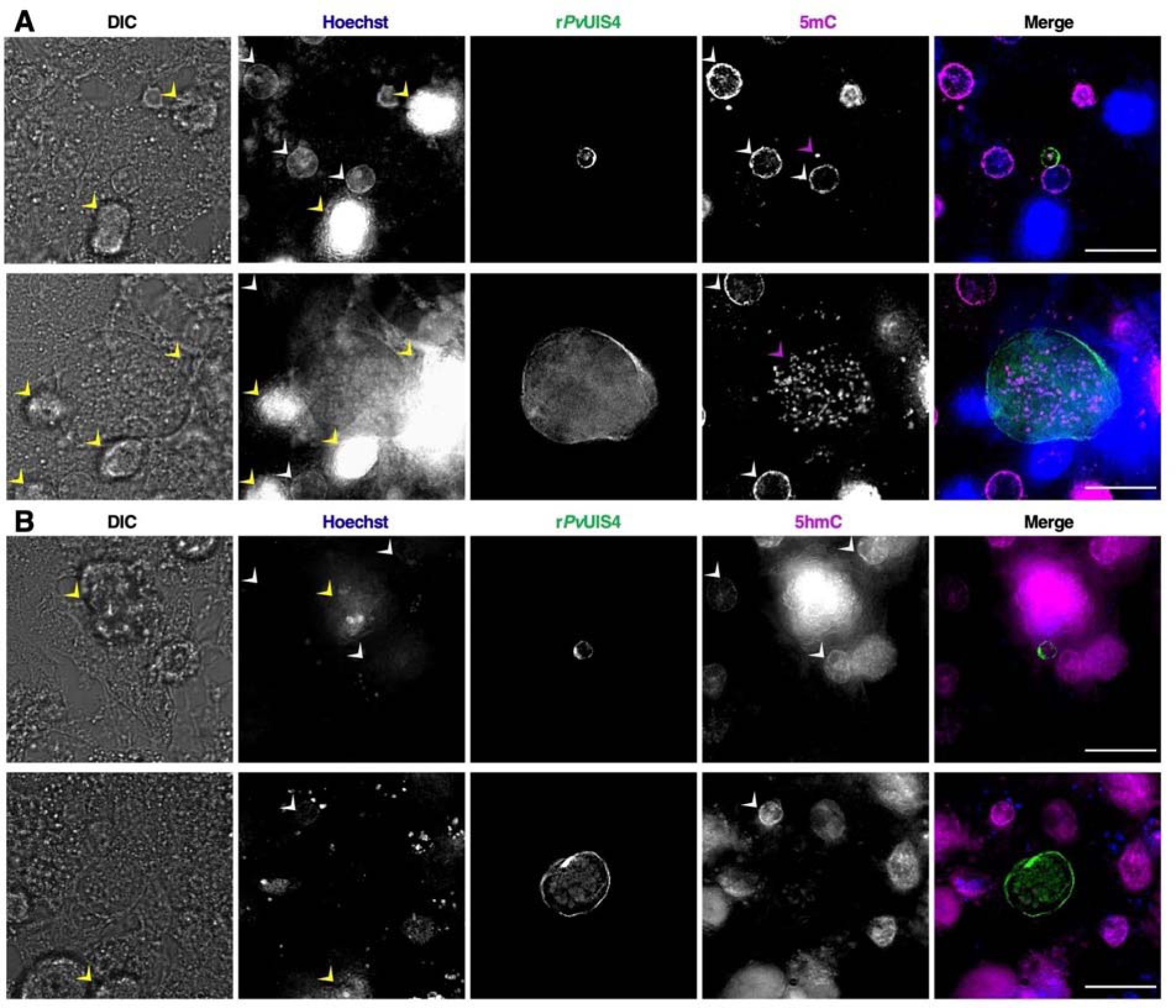
Cytosine modifications in *P. vivax* liver forms, full panels from Case 2. (**A**) Immunofluorescent imaging of a 5mC-positive *P. vivax* hypnozoite (top) and schizont (bottom) at day 6 post-infection. (**B**) Immunofluorescent imaging of a 5hmC-negative *P. vivax* hypnozoite (top) and schizont (bottom) at day 7 post-infection. Yellow arrows indicate autofluorescence in the blue channel associated with cell debris above the hepatocyte monolayer. White arrows indicate hepatocyte nuclei which are dimly-stained with Hoechst 33342 and positive for 5mC or 5hmC. Purple arrows indicate 5mC-positive foci within the parasite. Bars represent 20 µm.

**Fig. S8.**
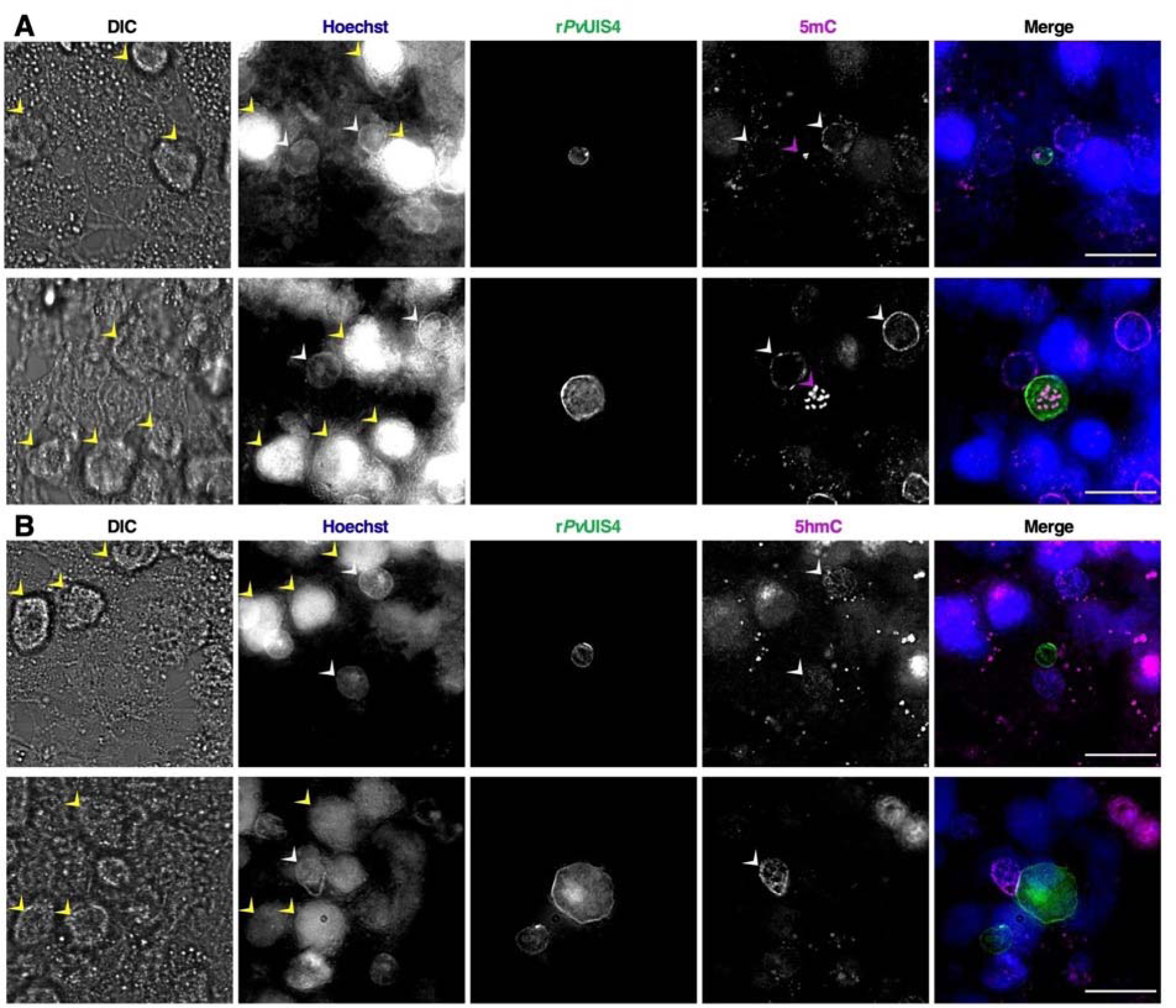
Cytosine modifications in *P. vivax* liver forms, full panels from Case 3. (**A**) Immunofluorescent imaging of a 5mC-positive *P. vivax* hypnozoite (top) and schizont (bottom) at day 6 post-infection. (**B**) Immunofluorescent imaging of a 5hmC-negative *P. vivax* hypnozoite (top) and schizont (bottom) at day 7 post-infection. Yellow arrows indicate autofluorescence in the blue channel associated with cell debris above the hepatocyte monolayer. White arrows indicate hepatocyte nuclei which are dimly-stained with Hoechst 33342 and positive for 5mC or 5hmC. Purple arrows indicate 5mC-positive foci within the parasite. Bars represent 20 µm.

**Fig. S9.**
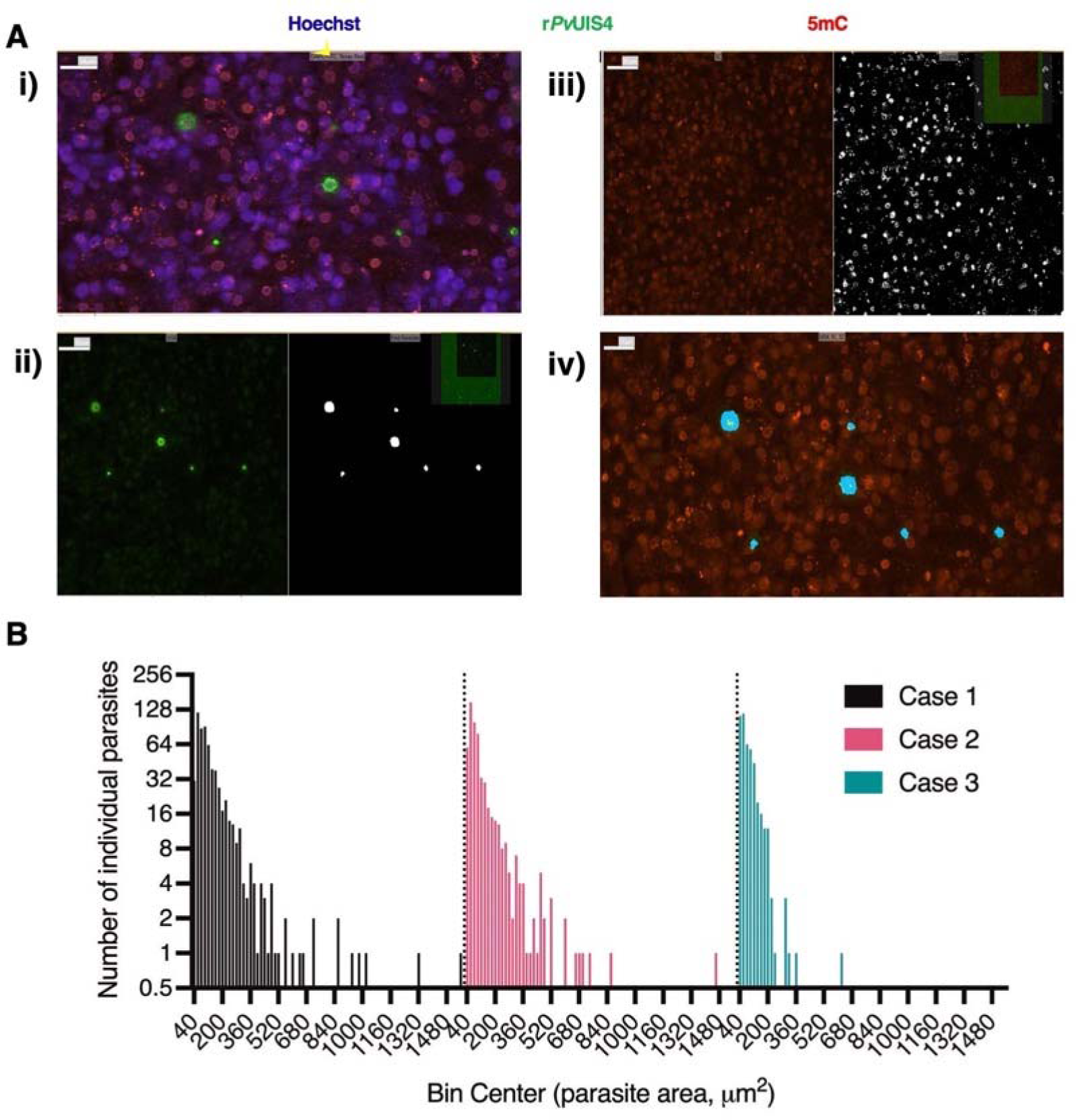
High-content analysis of cytosine modifications and *P. vivax* liver stage population metrics. (**A**) Masks used to quantify parasite area and 5mC or 5hmC signal, i) raw image taken with a 20x objective, ii) mask for *P. vivax* liver stages, iii) mask for 5mC or 5hmC signal, iv) intersection of parasite mask (light blue) and 5mC or 5hmC signal mask (yellow), leading to quantified area of signal per form. **(B**) Histogram of growth area all parasites quantified for Case 1, 2, and 3 in Fig. 3. Hypnozoites were classified as forms with an area 125µm^2^ and smaller.

**Fig. S10.**
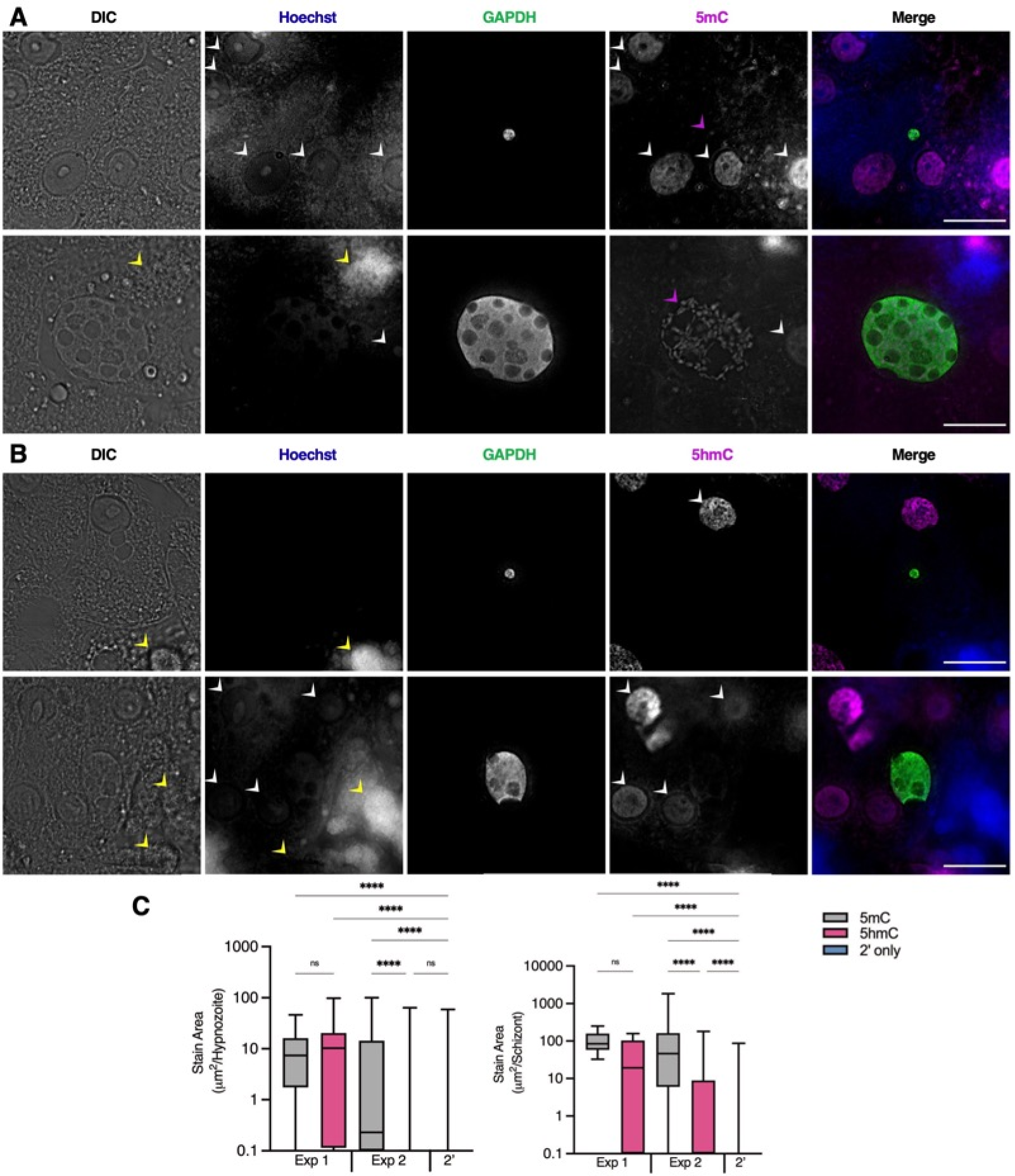
Cytosine modifications in P. cynomolgi M/B strain liver forms. (**A**) Immunofluorescent imaging of a 5mC-positive P. cynomolgi hypnozoite (top) and schizont (bottom) at day 8 post-infection. (**B**) Immunofluorescent imaging of a 5hmC-negative P. cynomolgi hypnozoite (top) and schizont (bottom) at day 8 post-infection. Yellow arrows indicate autofluorescence in the blue channel associated with cell debris above the hepatocyte monolayer. White arrows indicate hepatocyte nuclei which are dimly-stained with Hoechst 33342 and positive for 5mC or 5hmC. Purple arrows indicate 5mC-positive foci within the parasite. Bars represent 20µm. (**C**) High-content quantification of 5mC or 5hmC stain area within hypnozoites or schizonts. Experiment 1 was fixed at day 8 post-infection, Experiment 2 was fixed at day 12 post-infection. Significance determined using Kruskal-Wallis tests for hypnozoites and schizonts, with Dunn’s multiple comparisons, ****p <.0001, ns = not significant. Line, box and whiskers represent median, upper and lower quartiles, and minimum-to-maximum values, respectively, of all hypnozoites (124 ≤ n ≤ 712) or all schizonts (7 ≤ n ≤ 581) in culture, 2’ indicates a secondary stain only control. Images in A and B are from Experiment 1.

**Fig. S11.**
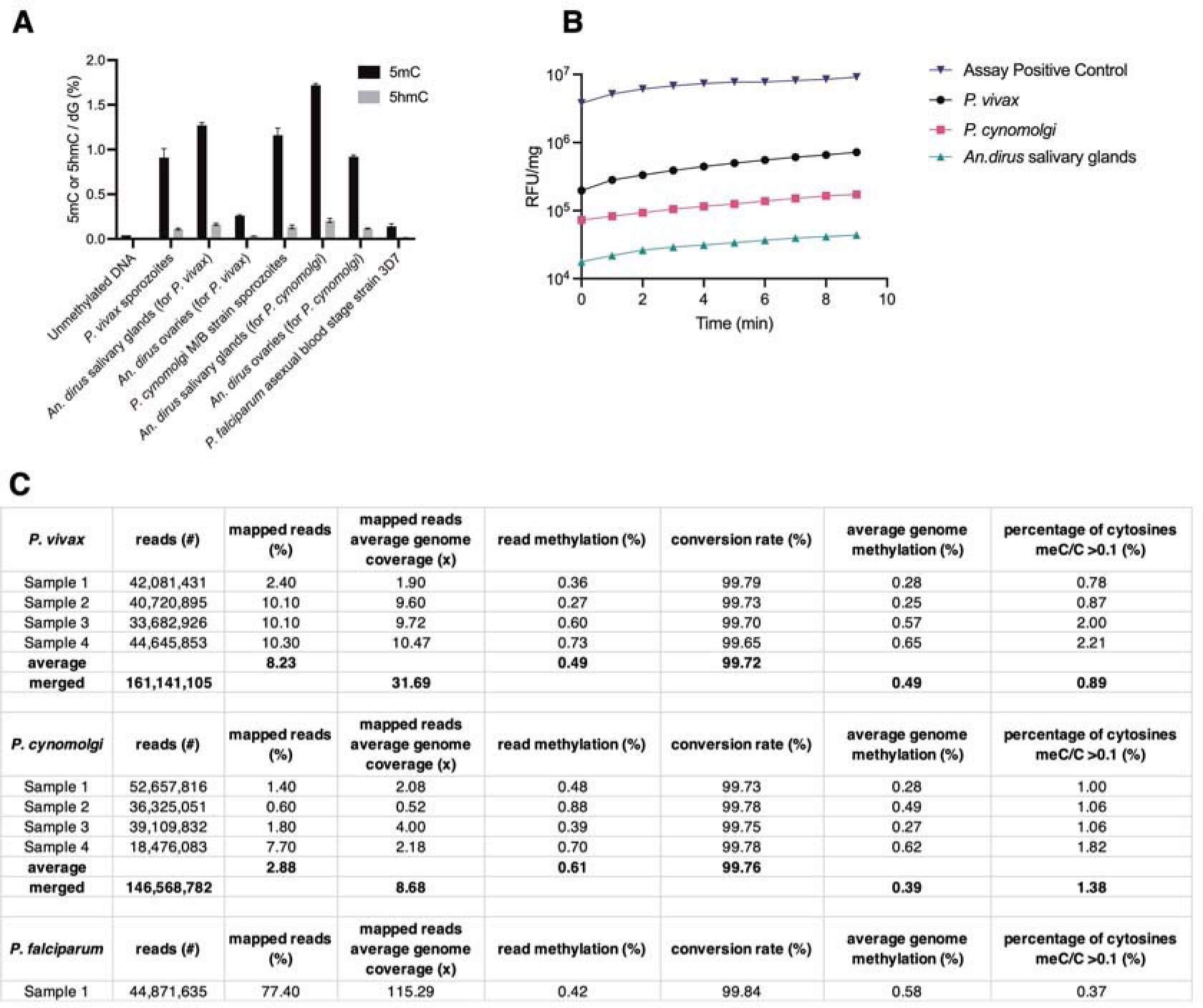
Measurement of DNA methylation and DNA methyltransferase (DNMT) in *P. vivax* and *P. cynomolgi* sporozoites. (**A**) Liquid chromatography-tandem mass spectrometry (LC-MS/MS) analysis of 5mC or 5hmC from enzymatically-digested gDNA from *P. vivax* sporozoites, *P. cynomolgi* sporozoites, and *P. falciparum* blood stage parasites, as well as negative controls including uninfected mosquito salivary glands and ovaries from the same colony of mosquitoes used to generate the respective sporozoites. Bars represent SD of two independent experiments. (**B**) DNMT activity measured from nuclear extracts of *P. vivax* sporozoites, *P. cynomolgi* sporozoites, and uninfected mosquito salivary glands using the Epiquick DNMT activity assay. Data are from a single experiment. (**C**) Summary statistics of read sets, percentage of mapped reads, read methylation levels, conversion rate and genome-wide methylation levels from bisulfite sequencing.

**Fig. S12.**
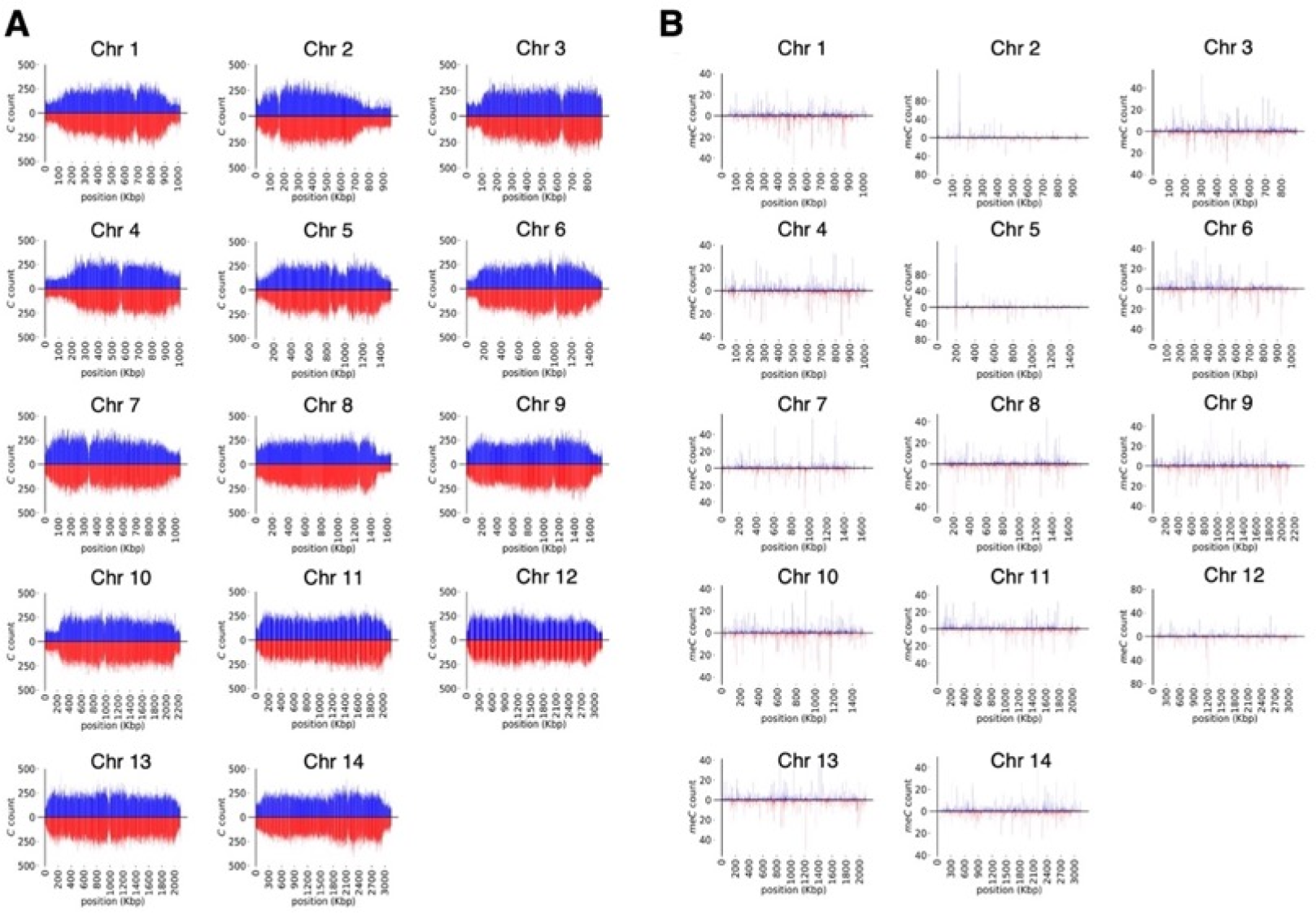
Cytosine and methylation density plots for *P. vivax* sporozoites. (**A**) CG content of chromosome 1 to 14 (Chr 1-14). The total number of cytosines quantified on each strand using 1 kbp long non-overlapping windows. (**B**) The total number of methylated cytosines quantified on each strand using 1 kbp long non-overlapping windows.

**Fig. S13.**
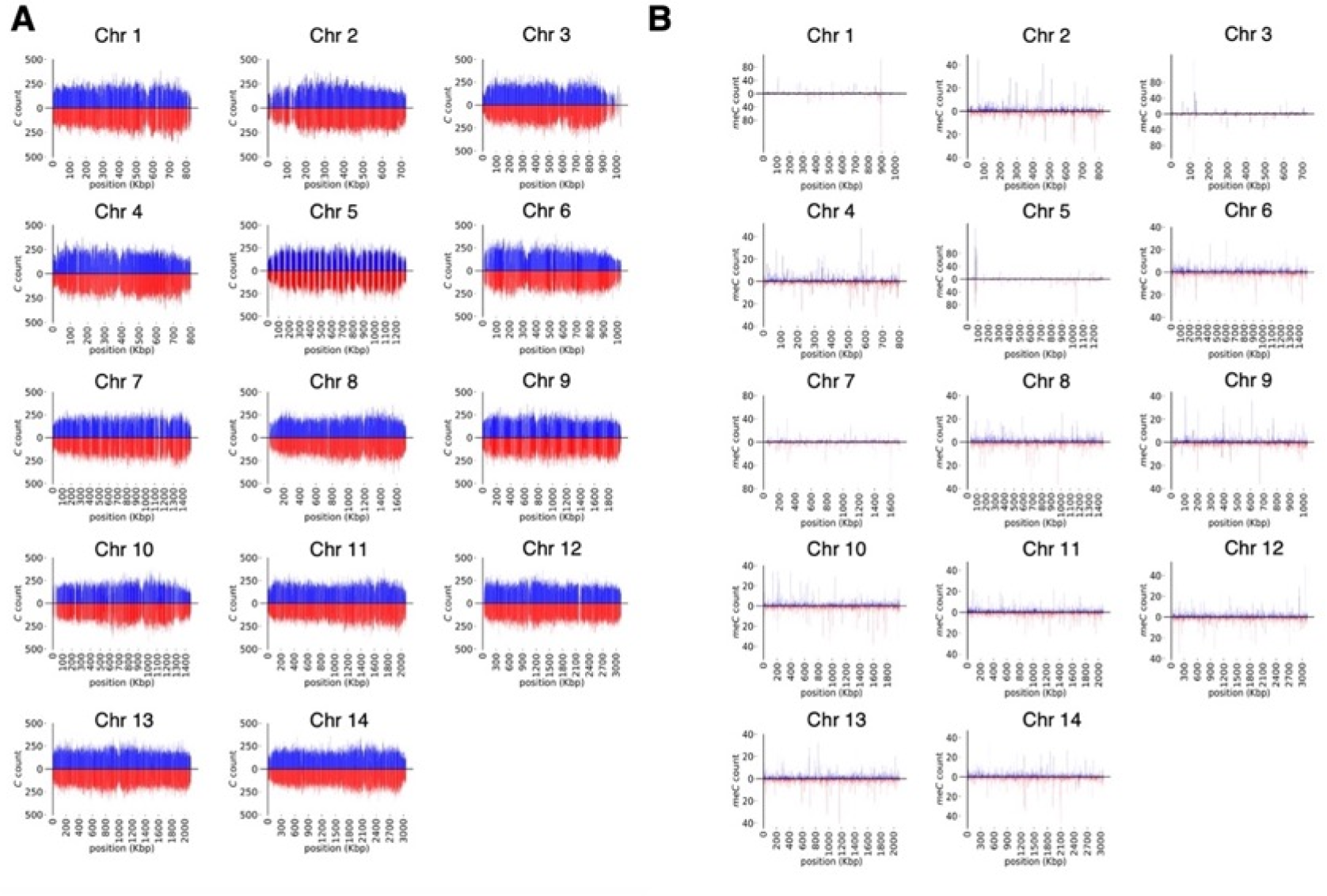
Cytosine and methylation density plots for *P. cynomolgi* sporozoites. (**A**) CG content of chromosome 1 to 14 (Chr 1-14). The total number of cytosines quantified on each strand using 1 kbp long non-overlapping windows. (**B**) The total number of methylated cytosines quantified on each strand using 1 kbp long non-overlapping windows.

**Fig. S14.**
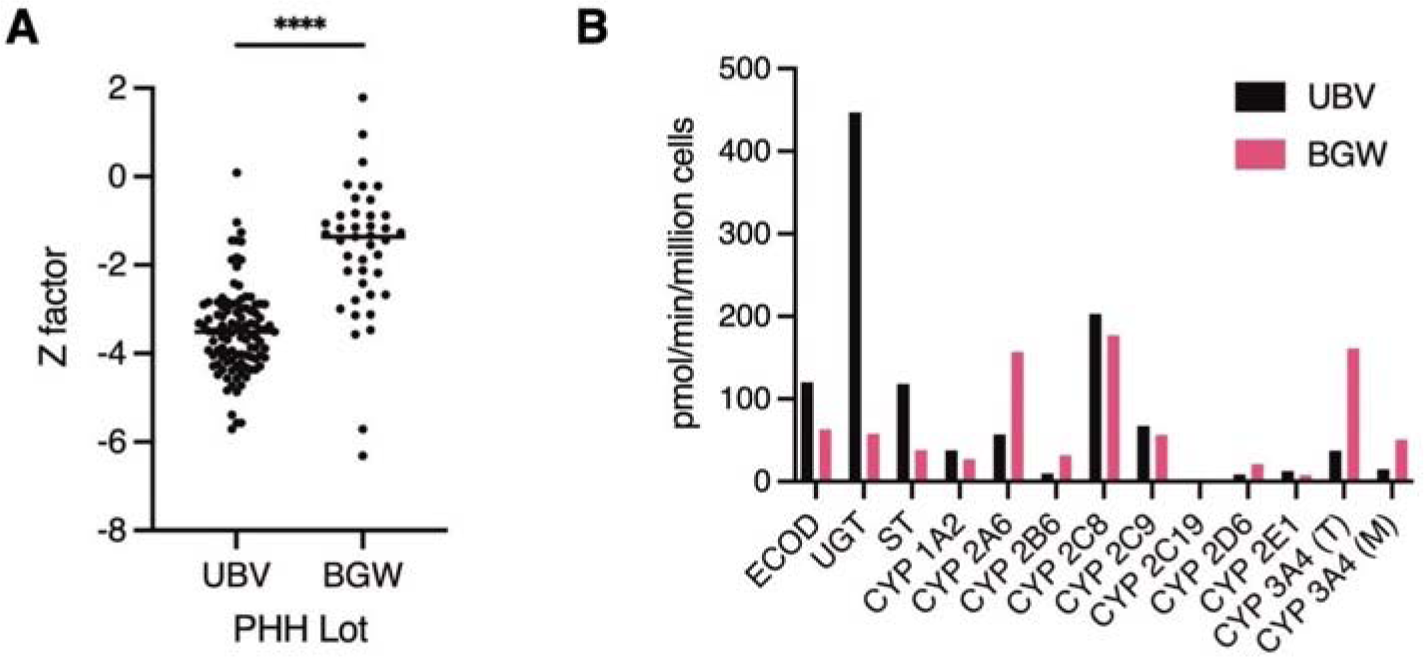
Monensin activity in all control wells based on PHH lot. (**A**) Initially, the ReFRAME was screened with cryopreserved vials of a specific lot of PHH, UBV. Screening continued with a new lot, BGW, once the supply of UBV vials was exhausted. The activity of monensin was significantly reduced in wells with BGW versus UBV PHH. A Mann-Whitney test indicates the difference was statistically significant, *U*(*N*_UBV_=105, *N*_BGW_=41) = 552, *z* = −2.15, \*\*\*\**p* <.0001. (**B**) Metabolic activity panel for PHH lots UBV and BGW performed as part of regular quality-control at the vendor (BioIVT). ECOD: 7-ethoxycoumarin O-deethylation, UGT: 7-hydroxycoumarin glucuronidation, ST: 7-hydroxycoumarin sulfation, CYP 1A2: phenacetin O-deethylation, CYP 2A6: coumarin 7-hydroxylation, CYP 2B6: bupropion hydroxylation, CYP 2C8: amodiaquine N-desethylation, CYP 2C9: tolbutamide methyl-hydroxylation, CYP 2C19: S-mephenytoin 4’-hydroxylation, CYP 2D6: dextromethorphan O-demethylation, CYP 2E1: chlorzoxazone 6-hydroxylation, CYP 3A4 (T): testosterone 6β-hydroxylation, CYP 3A4 (M): midazolam 1-hydroxylation.

**Fig. S15.**
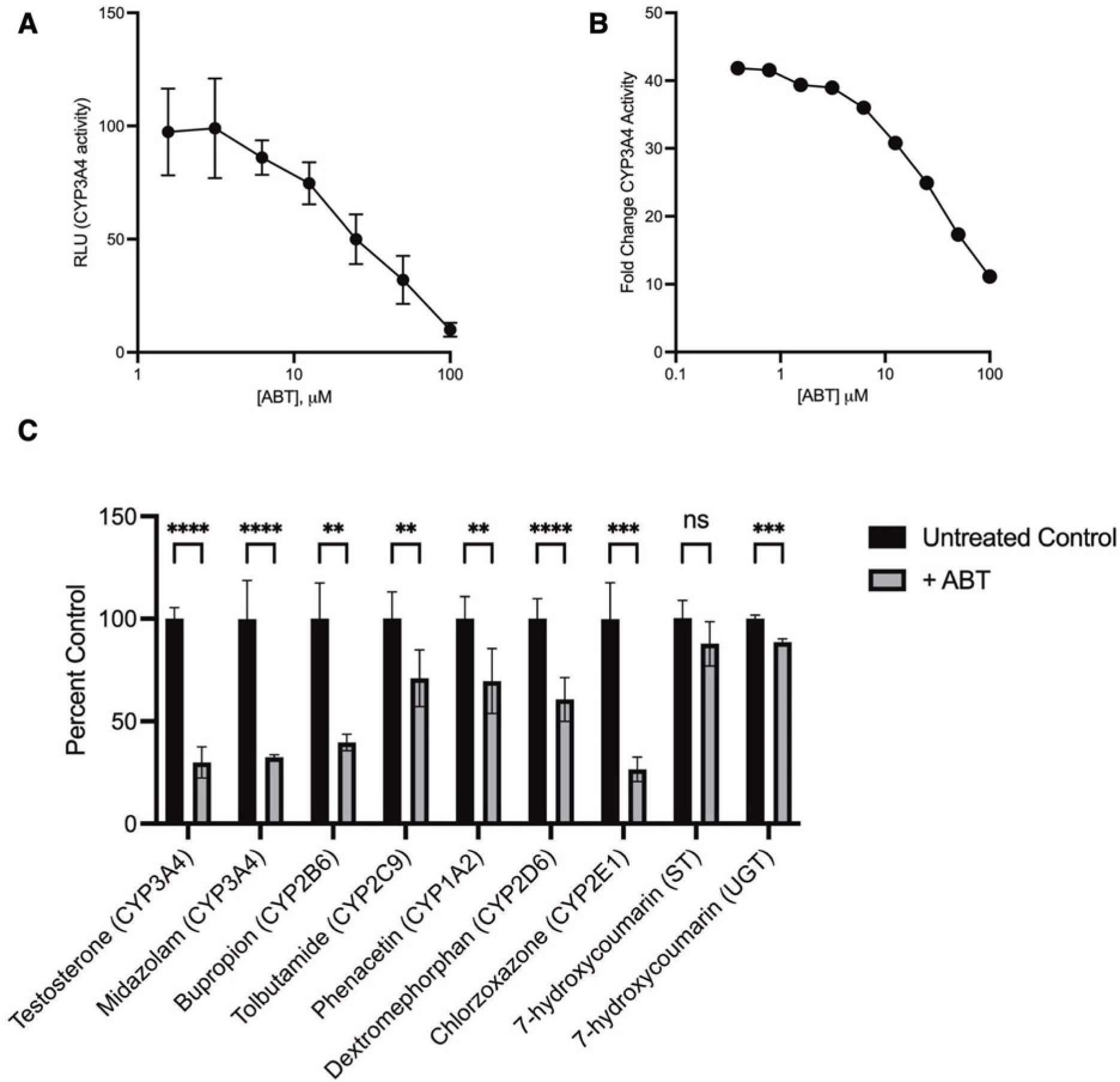
Characterization of primary human hepatocyte (PHH) metabolism following 1-aminobenzotriazole (1-ABT) treatment. (**A**) PHH lot BGW were seeded into 384-well plates and cultured for 7 days before addition of a dilution series of 1-ABT in media. Cytochrome P450 3A4 activity (CYP3A4) was measured using luciferin-IPA (Promega). RLU: relative luminescence units. Bars represent SD of quadruplicate wells. Data are representative of two independent experiments. (**B**) PHH lot BGW were cultured in 384-well plates before addition of 25 μM rifampicin in media on days 4 and 6 to induce CYP3A4 expression. At day 7 post-seed, CYP3A4 activity was measured by adding luciferin-IPA and a dilution series of 1-ABT in media. Fold change was calculated based on matching uninduced controls. Data are from one independent experiment. (**C**) PHH lot BGW were seeded in 384-well plates and cultured for 7 days before treatment with 100 μM 1-ABT for 1h, followed by addition of substrates for 1 h and collection for analysis by mass spectrometry. Data are combined from two independent experiments, bars represent SD of all replicates. Significance determined by student’s t tests, \*\*\*\**p* <.0001, \*\*\**p* <.001, \*\**p* <.01, ns, not significant.

**Fig. S16.**
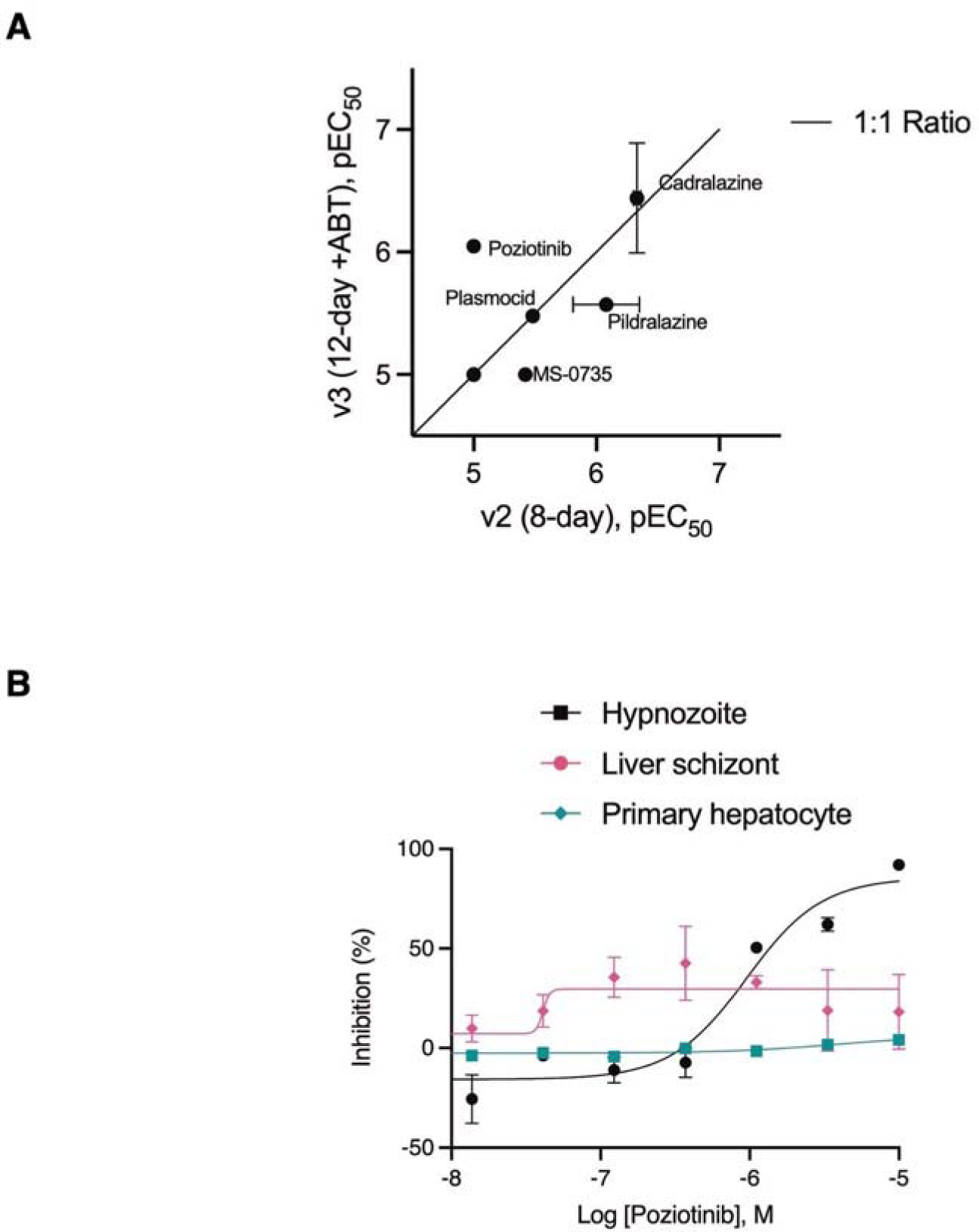
ReFRAME hits re-confirmed in a *P. vivax* 12-day 1-ABT assay. (**A**) Hypnozonticidal potency comparison of 12 ReFRAME hits in 8-day and 12-day 1-ABT dose-response confirmation assays. Cadralazine, plasmocid, and pidralazine potencies were unaffected by assay version, while MS-0735 was less potent, and poziotinib was more potent, in the 12-day 1-ABT assay. Budralazine, dramedilol, RGH-5526, dihydralazine, todralazine, endralazine, and mopidralazine were inactive (pEC50 < 5) regardless of assay version. (**B**) Dose-response chart of poziotinib activity in the 12-day 1-ABT assay, pEC_50_ against hypnozoites = 6.05. Bars represent SEM.

**Fig. S17.**
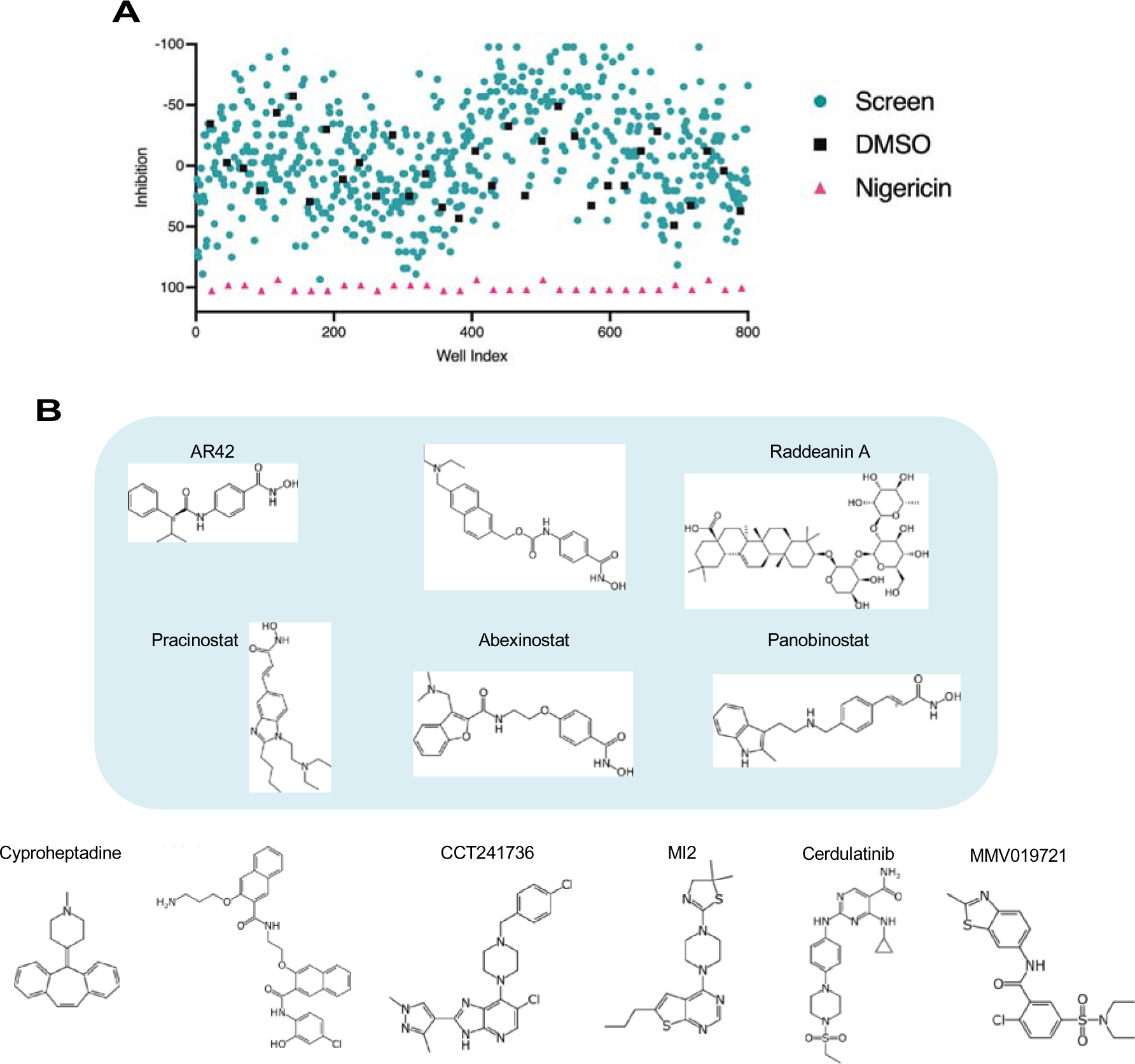
Epigenetic Inhibitor library screen and hits. (**A**) Index chart of an epigenetic inhibitor library screened against *P. vivax* hypnozoites in a v3 (12-day 1-ABT) assay. Teal circle: library, black square: DMSO, pink triangle: 200 nM nigericin. (**B**) Structures of epigenetic inhibitor hits which were confirmed to be active against *P. vivax* hypnozoites in dose-response assays; blue: histone deacetylase inhibitors.

**Fig. S18.**
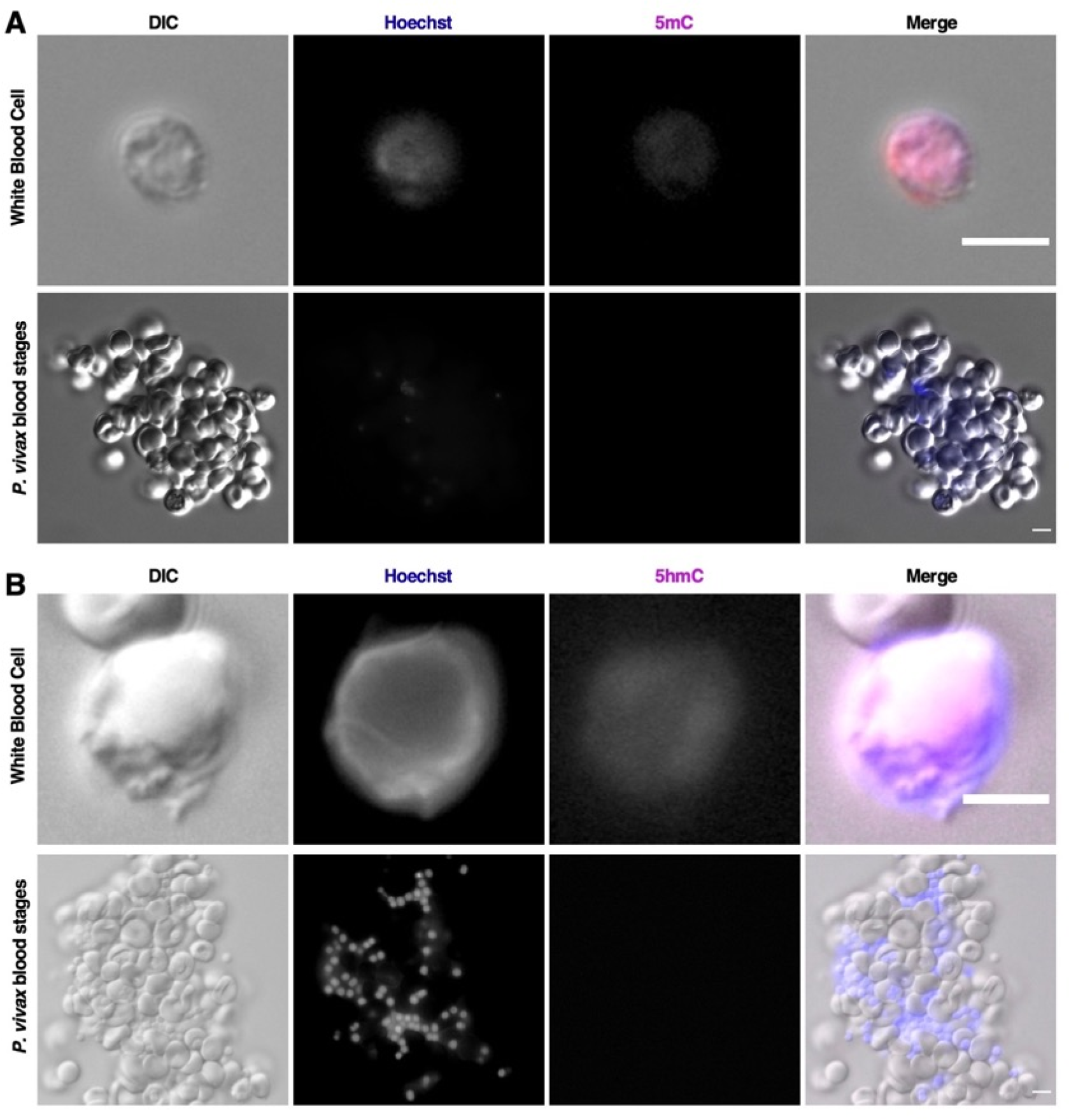
Cytosine modification in *P. vivax* blood stages. (**A**) *P. vivax* blood stages from patient isolates appeared negative when stained with 5mC. A white blood cell positive for 5mC serves as a stain control. (**B**) *P. vivax* blood stages from patient isolates appeared negative when stained with 5hmC. A white blood cell positive for 5hmC serves as a stain control. Bars represent 10 µm.

**Data S1.** (separate file)

Details for chemistry (batches, suppliers, structures); individual run data used to calculate all pEC_50_s and pCC_50_s; details for biologicals; details for reagents and replication.

**Data S2.** (separate file)

Raw pharmacokinetic data report from Wuxi for cadralazine in rhesus macaques.

**Data S3.** (separate file)

Contents of the Targetmol Epigenetic Library

## Notes

https://www.ncbi.nlm.nih.gov/sra/?term=PRJNA925570

### Summary of Updates

This version has been reorganized and now includes an additional synergy figure (Fig 2) and potency data for additional inhibitors (Table 1)

